# Development and application of aerobic, chemically defined media for *Dysgonomonas*

**DOI:** 10.1101/2020.08.11.247353

**Authors:** Charles M. Bridges, Daniel J. Gage

## Abstract

Members of *Dysgonomonas* are Gram-negative, non-motile, facultatively anaerobic coccobacilli originally described in relation to their isolation stool and wounds of human patients (CDC group DF-3). More recently *Dysgonomonas* have been found to be widely distributed in terrestrial environments and are particularly enriched in insect systems. Their prevalence in xylophagous insects such as termites and wood-feeding cockroaches, as well as in soil-fed microbial fuel cells, elicit interest in lignocellulose degradation and biofuel production, respectively. Their prevalence in mosquito and fruit fly have implications relating to symbiosis, host immunology and developmental biology. Additionally, their prevalence in termite, mosquito and nematode present novel opportunities for pest and vector control. Currently, the absolute growth requirements of *Dysgonomonas* are unknown, and they are cultured solely under anaerobic conditions on complex media containing blood, peptones, tryptones, and yeast, plant or meat extracts. Restrictive & undefined culturing conditions preclude physiological and genetic studies, and thus further understanding of metabolic potential. Here we describe the requirements for growth of termite-derived *Dysgonomonas* isolates and create parallel defined, minimal and complex media that permit vigorous and reliable aerobic growth. Furthermore, we show that these media can be used to easily enrich for *Dysgonomonas* isolates from complex and microbially-diverse environmental samples.

**Impact Statement:** Members of the genus *Dysgonomonas* are increasingly prevalent in ecological, medical and biotechnological contexts. To the best of our knowledge, there are currently no formulations for chemically defined or minimal media for *Dysgonomonas*, and limited complex formulations that allow aerobic growth, particularly on solid media. We have created three parallel media formulations (complex, defined & minimal) that permit robust aerobic and anaerobic growth in liquid and agar-solidified media. These formulations remove the necessity for animal blood and expensive equipment such as anaerobic chambers, which can inhibit basic research by groups with biosafety and resource limitations.

## Introduction

Members of the genus *Dysgonomonas* (family *Dysgonomonadaceae* [1]) once represented an emerging class of opportunistic pathogens isolated from human sources including stool, abscesses and wounds [2–6]. However, the majority of newly cultivated isolates originate from non-human environmental sources, including a soil-seeded microbial fuel cell [7], higher [8] and lower termites [9, 10], marine beach sand [11], paper mill sludge [12] and omnivorous cockroaches [13]. Sequences of the 16S rRNA gene of *Dysgonomonas* are prevalent in insect associations, such as honeybee [14], dipteran flies [15–18], including several *Drosophila* species [19–21], beetles [22–26], and several life stages of three major mosquito genera, *Culex* [27, 28], *Aedes* [29, 30] and *Anopheles* [31, 32]. *Dysgonomonas*-derived 16S rRNA gene sequences are routinely observed associated with xylophagous cockroaches [33–38] and as ectosymbionts of nematodes living in the cockroach digestive system [39], as well as in the hindguts of phylogenetically related termites [40–44]. *Dysgonomonas* have been identified as core microbiota of both higher [45] and lower [46] termites, including *Reticulitermes flavipes* [47]. There is growing interest in *Dysgonomonas* due to their prevalence in biotechnological processes such as lignocellulose degradation and bioconversion of polysaccharides for biofuel development [48, 49], microbial fuel and electrolysis cells [50–57], wastewater bioreactors [58, 59], biodegradation of food waste [60, 61] and pharmaceutical compounds [58, 62–64].

Members of genus *Dysgonomonas* are described as fastidious [3, 4, 65] owing to growth requirements satisfied only by rich, complex media containing whole or digested animal-derived components. For example, both the American Type Culture Collection (ATCC) and the German Collection of Microorganisms and Cell Cultures (DSMZ) recommend culture of *Dysgonomona*s on complex media containing enzymatic digests of animal-derived proteins, yeast- or animal-tissue extracts, defibrinated animal blood and other rarely required growth factors used by other fastidious organisms. Pre-mixed media formulations for these recommended media are widely available and can offer convenience, but are relatively expensive and cannot be easily tailored for experimental purposes. In-house preparations of these media can be made, but since many common recipes were created for the purpose of growing bacteria with diverse nutritional requirements, nearly all available media contain components that can be expensive or difficult to prepare and may be superfluous for the growth of *Dysgonomonas*. The Known Media Database (KOMODO) [66] is a collection of known organism-media pairings, and its online tool GROWREC (http://komodo.modelseed.org/growrec.htm) uses phylogenetic and ecological similarity to predict growth-permitting media for an organism. While this database performs well to determine complex anaerobic media formulations specific to *Dysgonomonas*, it is unable to offer either chemically defined or aerobic media formulations. Defined media created for the closely related genus *Bacteroides* [67, 68] have been shown to support the growth *Dysgonomonas gadei* and *Dysgonomonas mossii* [69], but we observed poor growth from our isolates in these liquid media when serially cultured (unpublished results). *Dysgonomonas alginatilytica* has been reported to grow on a defined medium consisting only of basal salts and kraft-lignin [12], and while this medium may contain adequate reduced carbon and concomitant micronutrients, it lacks major components of the media known to support the growth of *Dysgonomonas*. Defined media have been described for closely-related *Porphyromonas gingivalis* [70, 71], but contain components such as casitone that are not truly defined, or lack known required nutritional components such as B-vitamins. Described species of *Dysgonomonas* are facultatively anaerobic on complex media [3–5, 7–9, 11, 72], but our isolates exhibited slow growth and altered colony morphology when grown aerobically on available complex media formulations. ATCC recommends that members of *Bacteroidaceae* be grown anaerobically using pre-reduced medium containing sodium sulfide, cysteine, or coenzyme M as reducing agents (ATCC Bacterial Culture Guide, 2015). These reductants, and expensive equipment such as anaerobic chambers, may be unnecessary given the facultative nature of *Dysgonomonas*.

The lack of chemically defined media for members of *Dysgonomona*s precludes a deeper understanding of physiological and metabolic potential and dampens the ability to perform phenotypic analyses. Furthermore, defined media complement the development of genetic tools, such as the addition or omission of specific components for the purpose of genetic selections. Here, we developed chemically defined media formulations based on the minimal growth requirements of four *Dysgonomonas* isolates belonging to differing phylogenetic clades within the genus. The parallel media formulations *Dysgonomonas* <u>Complex</u> Medium (D<u>C</u>M), *Dysgonomonas* <u>Defined</u> Medium (D<u>D</u>M) and *Dysgonomonas* <u>Minimal</u> Medium (D<u>M</u>M) make use of widely available components, offer straightforward preparation, allow growth in liquid or solid culture and permit growth under aerobic or anaerobic conditions. Finally, we demonstrate the ease with which our minimal media (DMM) allows enrichment of *Dysgonomonas* from microbially-complex environmental samples such as the lower termite hindgut.

## Materials and methods

### Isolation & maintenance of bacteria from termite hindguts

*Dysgonomonas* spp. BGC7 and HGC4 were generously provided by Dr. Michael C. Nelson and Dr. Joerg Graf at the University of Connecticut. *R. flavipes* termites were collected from decaying oak tree stumps in Granby, CT, USA (Lat: 41.999316, Long: −72.789053) during September 2017. All isolations were performed in a laminar flow hood using sterile instruments and reagents. *Dysgonomonas* spp. GY75 and GY617 were isolated by washing ten worker termites in 95% ethanol and aseptically extirpating hindguts using the technique described by Matson et al. [73]. Hindgut contents were pooled into 1 ml sterile 1X M9 salts (5.8 g/L Na_2_HPO_4_ (Fisher, Pittsburgh, PA, USA), 3.0 g/L KH_2_PO_4_ (Fisher), 0.50 g/L NaCl (Fisher), 1.0 g/L NH_4_Cl (Fisher), 0.011 g/L CaCl_2_ (Acros Organics, Geel, Belgium), 0.25 g/L MgSO_4_ (Acros)) and the entire volume added to a 2-ml screw-capped microcentrifuge tube (USA Scientific, Ocala, FL, USA) containing 200 mg of 1 mm glass beads (Biospec, Bartlesville, OK, USA) and processed for 30 seconds on a Mini-Beadbeater-16 (Biospec). Homogenized samples were serially diluted in sterile 1X M9 salts and plated onto peptone-yeast extract-blood-glucose medium (PYBG; 6 g/L Na_2_HPO_4_·7H_2_O, 10 g/L proteose peptone #3 (US Biological, Salem, MA, USA), 10 g/L yeast extract (Bacto, Mt Pritchard, NSW, Australia), 10% whole sheep blood (Lampire, Pipersville, PA, USA), 0.5% D-glucose (Acros), 50 μg/ml kanamycin sulfate (Gibco, Dublin, Ireland) and 1.5% w/v agar (Bacto)). Plates were incubated anaerobically for 5 days at 22°C in a gloveless anaerobic chamber (Coy, Grass Lake, MI, USA) under an atmosphere of 5.5% CO_2_, 5.5% H_2_ and 89% N_2_. Isolates with colony morphology similar to *Dysgonomonas* to were serially sub-streaked twice onto the same medium and incubated under the same conditions. Purified isolates of two strains, GY75 and GY617, were grown under anaerobic conditions at 22°C in rich peptone-hemin-glucose medium (rPHG; 6 g/L Na_2_HPO_4_·7H_2_O, 30 g/L proteose peptone #3, 10 g/L yeast extract, 50 mg/L porcine ferric hemin (MP Biomedicals, Santa Ana, CA, USA; prepared as an aqueous solution of 0.5 g/L in 10 mM NaOH (Fisher)), 0.5% w/v D-glucose, 50 μg/ml kanamycin sulfate, 1% v/v Wolfe’s Mineral Solution [74] (WMS; 3.0 g/L MgSO_4_·7H_2_O (Acros), 1.5 g/L nitrilotriacetic acid (Acros), 1.0 g/L NaCl, 0.5 g/L MnSO_4_·2H_2_O (Acros), 0.1 g/L CoCl_2_·6H_2_O (Sigma, St. Louis, MO, USA), 0.1 g/L ZnSO_4_·7H_2_O (Research Organics, Cleveland, OH, USA), 0.1 g/L CaCl_2_·2H_2_O (Acros), 0.1 g/L FeSO_4_·7H_2_O (Research Organics), 0.025 g/L NiCl_2_·6H_2_O (Acros), 0.02 g/L KAl(SO_4_)_2_·12H_2_O (Fisher), 0.01 g/L CuSO_4_·5H_2_O (Sigma), 0.01 g/L H_3_BO_3_ (Fisher), 0.01 g/L Na_2_MoO_4_·2H_2_O (Sigma), 0.3 g/L Na_2_SeO_3_·5H_2_O (Sigma)) and 5% v/v Wolfe’s Vitamin Solution [74] (WVS; 2.0 mg/L biotin (Fisher), 2.0 mg/L folic acid (Sigma), 10 mg/L pyridoxine-HCl (Sigma), 5.0 mg/L thiamine-HCl (Sigma), 5.0 mg/L riboflavin (Sigma), 5.0 mg/L nicotinamide (Sigma), 5.0 mg/L calcium-D-pantothenate (Sigma), 100 μg/L cyanocobalamin (Fisher), 5.0 mg/L p-aminobenzoic acid (PABA; Sigma), 5 mg/L *α*-lipoic acid (Sigma)), pH 7.5) before being frozen at −80°C in 20% (v/v) sterile glycerol (Acros). Isolates were cultured form glycerol stocks and maintained on solid PYBG medium incubated anaerobically for 72 hours at 22°C. Unless otherwise stated, cultures used as growth curve inocula were grown anaerobically at 22°C to stationary phase in Falcon 15 ml polypropylene conical tubes (Corning, Corning, NY, USA) in 5 ml volumes.

### Identification of bacterial isolates

The identity of four strains of *Dysgonomonas* (isolates BGC7, HGC4, GY75 & GY617) were confirmed by sequencing nearly full-length 16S rRNA gene sequences (∼1455 of 1535 bp, hereafter called ‘full-length’). Single colonies from pure isolates were inoculated into 5 ml rPHG medium in 18 x 150 mm glass culture tubes (Fisher) containing 50 μg/ml kanamycin sulfate and incubated ∼18 hours at 30°C aerobically without shaking. Overnight cultures were used for genomic DNA (gDNA) preparation using Promega Wizard Genomic DNA Purification Kit (Promega, Madison, WI, USA) or Epicenter MasterPure Complete DNA and RNA Purification Kit (Lucigen, Madison, WI, USA) according to manufacturer instructions. Isolated gDNA was checked for quality and concentration by gel electrophoresis and by spectrophotometry using a Take3 Micro-Volume Plate (BioTek, Winooski, VT, USA). Approximately 100 ng of gDNA was used as template for PCR using Q5 Hot Start DNA Polymerase (New England Biolabs (NEB), Ipswich, MA, USA) and primers Eubact_27F (5’-AGAGTTTGATCMTGGCTCAG-3’) and Eubact_1492R (5’-TACGGYTACCTTGTTAC-3’) (modified from [75]). A no-template control contained water in place of gDNA. PCR was performed using the manufacturer’s recommended procedure with a primer annealing temperature of 50°C and a 45 second extension time, for a total of 35 cycles. An aliquot of each PCR reactions was checked by gel electrophoresis for single amplicons before purifying the remaining reaction using Monarch DNA Gel Extraction Kit (NEB), per manufacturer instructions. PCR amplicons were A-tailed by incubating 750 ng of DNA with 2 Units of Taq DNA Polymerase (NEB) and 0.2 mM dATP (Thermo Fisher Scientific, Inc., Waltham, MA, USA) at 72°C for 30 minutes. A-tailed PCR amplicons were cloned into p-GEM-T Easy Vector System (Promega) per manufacturer instructions and ligation reactions introduced directly into electrocompetent XL1-Blue *E. coli* (Stratagene, La Jolla, California, USA) by electroporation. After a 1 hour recovery at 37°C in liquid SOC (10 g/L tryptone (Bacto), 2.5 g/L yeast extract, 0.6 g/L NaCl, 0.2 g/L KCl (Fisher), 3.6 g/L D-glucose, 2 g/L MgCl_2_·6H_2_O (Research Organics), cells were diluted 10-fold and plated on agarose-solidified LB (10 g/L tryptone, 5 g/L yeast extract, 10 g/L NaCl, 15 g/L agar) containing 100 μg/ml ampicillin sodium salt (Fisher), 47.6 μg/ml (0.2 mM) isopropyl β-D-thiogalactopyranoside (IPTG; Fisher), 40 μg/ml (97.8 μM) 5-bromo-4-chloro-3-indolyl-β-D-galactopyranoside (X-gal; Fisher). Plates were incubated ∼18 hours at 37°C and individual white colonies screened for insert by colony PCR using primers M13_F (5’-ACGACGTTGTAAAACGACGGCCAGT-3’) and M13_R (5’-ATTTCACACAGGAAACAGCTATGACCA-3’) GoTaq DNA Polymerase (Promega). Reactions were separated by gel electrophoresis, and insert-positive clones were cultured ∼18 hours in 5 ml LB broth containing 100 μg/ml ampicillin. Plasmid DNA was isolated using Promega PureYield Plasmid Miniprep System (Promega) and sent to Genewiz (South Plainfield, NJ, USA) for bidirectional dideoxy sequencing using primers M13_F and M13_R. Overlapping sequencing reads were assembled and 2-4X sequence coverage was obtained for each clone.

### Growth experiments

Experiments were all performed in a basal media containing 1X M9 salts, and when required, amended by adding components to the following final concentrations (unless otherwise stated): 1% w/v proteose peptone #3, 1% w/v casamino acids (Fisher), 0.1 mg/ml L-tryptophan (Sigma), 0.1 mg/ml L-cysteine-HCl·H_2_O (Fisher), 0.1 mg/ml L-methionine (Sigma), 0.1 mg/ml each of nucleobases adenine, guanine, cytosine, thymine, uracil, hypoxanthine (Acros), 0.2 mg/ml L-ornithine-HCl (Acros), 0.2 mg/ml beta-nicotinamide adenine dinucleotide reduced disodium salt (NAD; Alfa Aesar, Lancashire, United Kingdom), 0.2 mg/ml diaminopimelic acid (DAP; Sigma), 1% v/v WMS, 5% v/v WVS, 0.5% w/v D-glucose, 50 μg/ml kanamycin sulfate, 10% v/v hemin solution. Unless otherwise stated, cell cultures used for growth experiments were pre-grown anaerobically without shaking for ∼18 hours in 5 ml volumes of rPHG medium. Cultures were centrifuged and the cell pellet washed three times with 1X M9 salts. Optical density of the cultures at 595 nm was determined by spectrophotometry using BioTek Synergy HT or H1 automated plate readers (BioTek) then diluted to an OD_595_ of 0.1 (200 μl of 1X M9 salts in 96-well microtiter plate (Corning #35-1172)) before being added to media at a final OD_595_ of approximately 0.01 in 1X M9 salts. When stated, cells were starved of particular media components prior to performing growth curves as follows. Cultures pre-grown in rPHG medium were washed and diluted to OD_595_ of 0.1 as described above, and subcultured 1:100 into 50 ml volume of defined medium or minimal medium lacking the specific component(s). Cultures were grown to component-limiting stationary phase before being washed, diluted and used as inoculum as stated above. When the presence of sulfate or ammonium was of concern, cells were washed and resuspended in sterile 1X phosphate-buffered saline (PBS; 8.0 g/L NaCl, 0.20 g/L KCl, 1.44 g/L Na_2_HPO_4_, 0.24 g/L mM KH_2_PO_4_, pH 7.4) in place of 1X M9 salts. Water used to prepare all components and media was Fisher Optima HPLC-grade water (Fisher). Aerobic growth curves were performed using BioTek HT and H1 Synergy microplate readers, using 220 μl total volume in 96-well micro-titer plates at 30°C. Automated optical density readings at 595 nm (OD_595_) were taken every 10 minutes following a 30-second orbital shake. Where described, duplicate plates were inoculated and each plate grown in parallel under aerobic or anaerobic conditions as described above. Endpoint OD_595_ readings were taken at the start and finish of each experiment, preceded by a 2-minute orbital shake. Unless otherwise indicated, all experiments included 2 biological replicates per plate with the exception of (i) auxanography for amino acids (**Figures S1 & S2**) & vitamins (**Figure S7**) which were performed as preliminary screens and subsequently followed by experiments to validate observed phenotypes; (ii) experiments examining a broad range of component concentrations such as L-cysteine or ferric hemin (**Figures S3, S5 & S6**). All experiments regardless of biological replicates were performed multiple times. Generation of graphs, as well as regression and statistical analyses were performed using GraphPad Prism v8 (GraphPad Software, La Jolla, CA, USA).

### Amino acid auxanography

To reveal possible amino acid auxotrophies in wild-type *Dysgonomonas* isolates, a modified version of auxanography [76] was performed (**Figures S1 & S2**). To reduce experimental complexity, B-vitamins were supplied. Nucleobases, NAD and PABA were omitted, as they were not required for growth (**Figure 1**). Twenty-two individual amino acids (obtained from Sigma except for L-cysteine-HCl·H_2_O), were prepared as 10 mg/ml aqueous solutions using concentrated NaOH or HCl to allow dissolution as necessary. All were free acids with the exception of the following chloride salts: L-ornithine-HCl, L-arginine-HCl, L-histidine-HCl, L-lysine-HCl and L-cysteine-HCl·H_2_O. Equal volumes of each amino acid in a pool were combined to 6 mL total volume, with each amino acid at a final concentration of 1.67 mg/ml. Pools can be found in **Table S1** and as follows: Pool 1: L-phenylalanine, L-serine, L-tryptophan, L-tyrosine, L-glutamine; Pool 2: L-alanine, L-cysteine, L-threonine, L-asparagine, L-methionine, DAP; Pool 3: L-arginine, L-ornithine, L-aspartic acid, L-proline, L-glutamic acid; Pool 4: L-leucine, L-glycine, L-isoleucine, L-histidine, L-lysine, L-valine; Pool 5: L-phenylalanine, L-alanine, L-arginine, L-leucine; Pool 6: L-serine, L-cysteine, L-ornithine, L-glycine; Pool 7: L-tryptophan, L-threonine, L-aspartic acid, L-isoleucine; Pool 8: L-tyrosine, L-asparagine, L-proline, L-histidine; Pool 9: L-methionine, L-glutamic acid, L-lysine; Pool 10: L-glutamine, DAP, L-valine. Media were supplemented with amino acid pools at a ratio of 1:5 such that each amino acid was present at 0.33 mg/ml each. The final media consisted of 1X M9 supplemented with 5% v/v hemin solution, 5% WVS, 0.5% v/v WMS, 0.5% w/v D-glucose, 50 μg/ml kanamycin sulfate and a single amino acid pool. Cells were pre-grown in rPHG, washed and used as inoculum as described above, and growth curve analysis performed.

**Figure 1.**
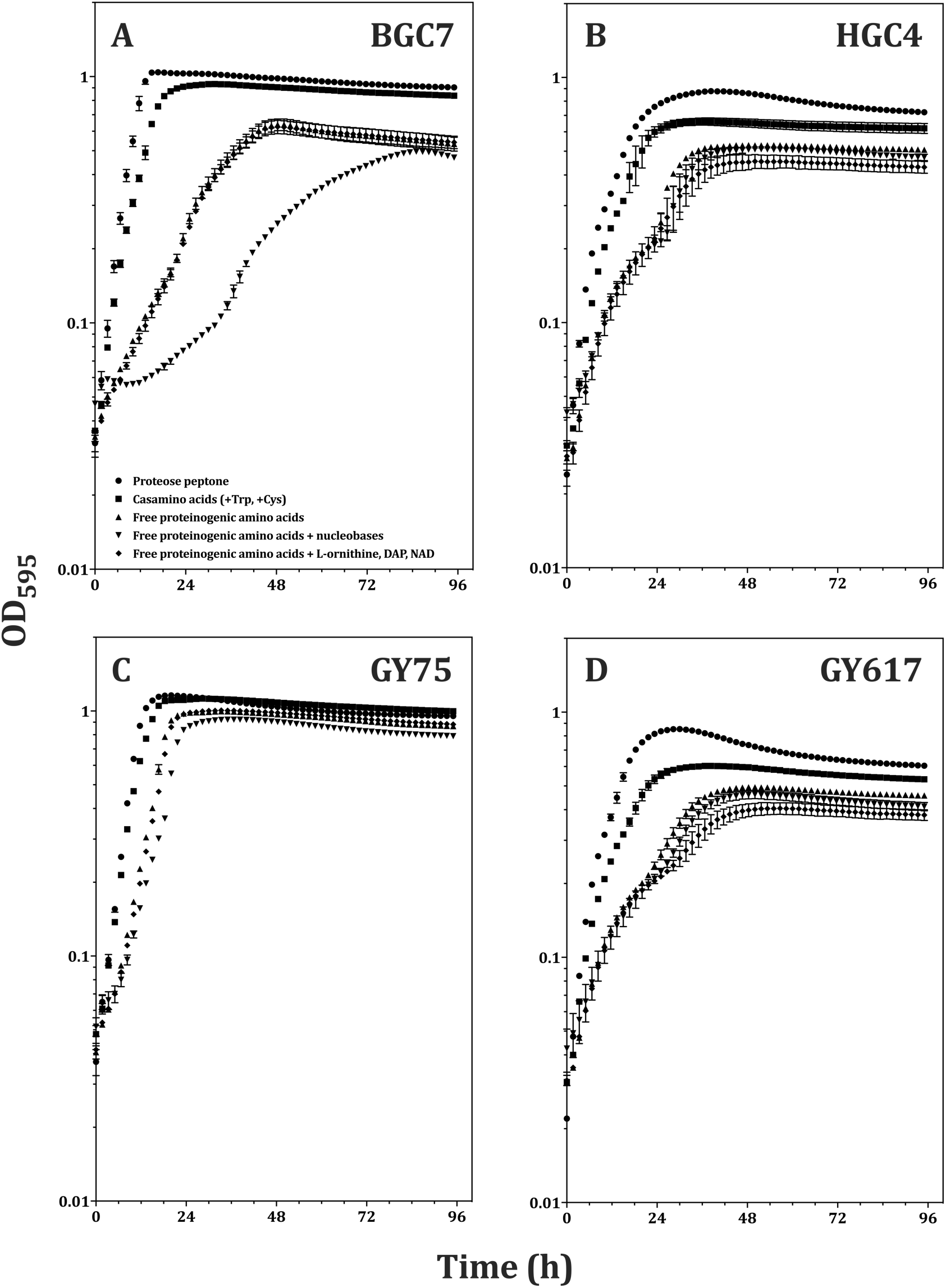
Growth of *Dysgonomonas* isolates in defined medium containing amino acids and B-vitamins. *Dysgonomonas* isolates BGC7 (A), HGC4 (B), GY75 (C) or GY617 (D) grown in 1X M9 salts supplemented with 5% v/v hemin solution, 5% v/v WVS, 0.5% v/v WMS 0.5% w/v D-glucose, 50 μg/ml kanamycin sulfate, pH 7.5. Amendments are shown in the legend in panel A. Error bars represent standard error of the mean (n=2).

### Determination of sulfur, nitrogen and iron requirements

To determine L-cysteine and sulfate requirements (**Figure 2**), cells were pre-grown in DDM containing 0.569 mM L-cysteine, 1 mM MgSO_4_, 5% v/v WVS and 10% v/v hemin solution before being washed and diluted in 1X PBS and used for growth curves. Modified WMS (mWMS) was created by substituting magnesium chloride or manganese chloride (MnCl_2_·4H_2_O, Acros) salts for the magnesium and manganese sulfate salts in the standard WMS recipe. Growth curves were performed in either sulfate-replete (0.01% v/v WMS, 1 mM total SO_4_^2-^) or sulfate-limited (0.01% v/v mWMS, 83 nM total SO_4_^2-^) conditions. To determine if L-cysteine was able to be utilized as a sole source of nitrogen (**Figure S4**), cultures were pre-grown in rPHG, washed and resuspended in PBS, and used as inoculum for growth curves in the presence or absence of excess ammonium (18.7 mM from 1X M9) and/or 1.7 mM L-cysteine. For determination of growth response to hemin concentration (**Figures S5 & S6**) and for determination of alternate sources of iron beside ferric hemin (**Figure 3**), cells were pre-grown in rPHG, then washed and subcultured 1:100 in 50 ml DDM without hemin and grown to hemin-limited stationary phase. Cells were again washed, diluted and then used for growth curves.

**Figure 2.**
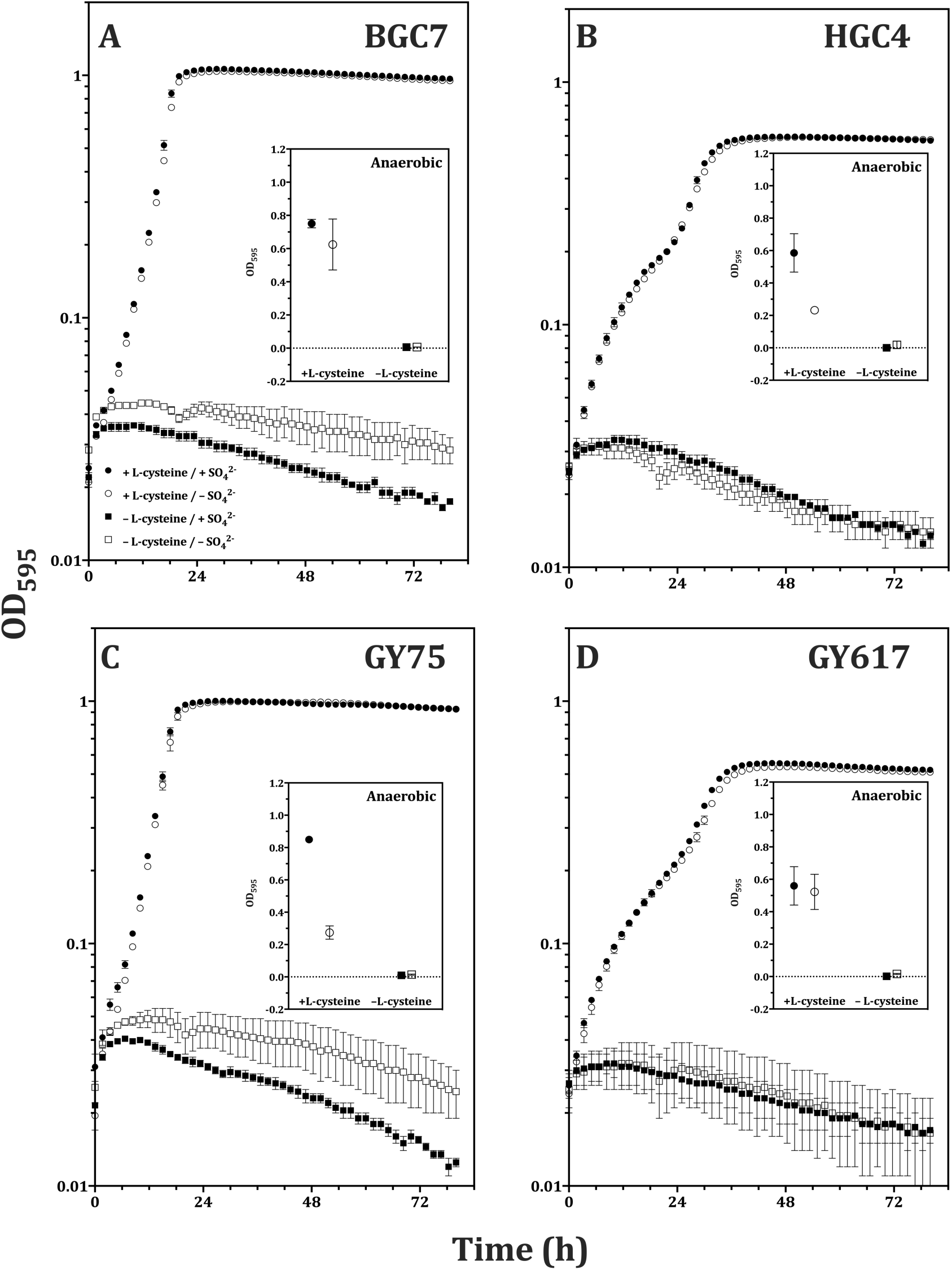
L-cysteine is required for aerobic and anaerobic growth. *Dysgonomonas* isolates BGC7 (A), HGC4 (B), GY75 (C) or GY617 (D) were pre-grown in DDM and diluted into sulfate-replete medium (1X M9 salts, 0.01% mWMS, 10% v/v hemin solution, 5% v/v WVS, 0.5% w/v D-glucose, pH 7.5; 1mM total sulfate) or sulfate-limited medium (1X M9 salts with 1mM MgCl_2_ replacing MgSO_4_, 0.01% WMS, 10% v/v hemin solution, 5% v/v WVS, 0.5% w/v D-glucose, pH 7.5; 83 nM total sulfate). Media contained either 0 or 1.7 mM L-cysteine as stated. Kanamycin sulfate was omitted for all conditions. Insets show final corrected OD_595_ under anaerobic conditions; the y-axis is in linear units. L-cysteine and sulfate combinations can be found in the legend in panel A. Error bars represent standard error of the mean (n=2)

**Figure 3.**
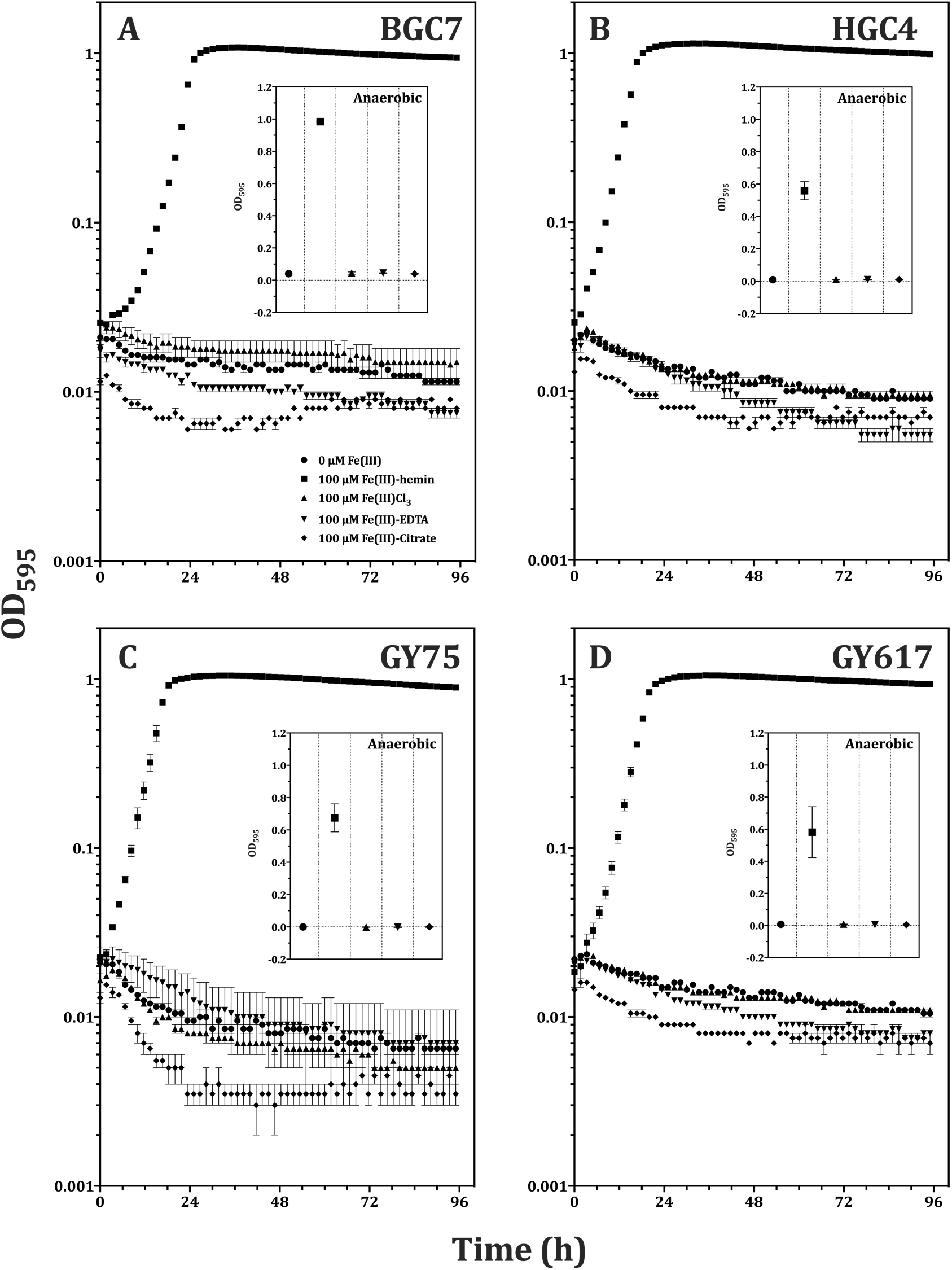
Ferric hemin is a preferred source of iron for *Dysgonomonas.* Isolates BGC7 (A), HGC4 (B), GY75 (C) or GY617 (D) were pre-grown in DDM washed and diluted into 1X M9 salts supplemented with 1% v/v WMS, 0.5% w/v D-glucose, 1.7 mM L-cysteine, 50 μg/ml kanamycin sulfate, pH 7.5, and contained iron sources according to the legend in panel A. Insets show final corrected OD_595_ under anaerobic conditions; the y-axis is in linear units. Error bars represent standard error of the mean (n=2).

### Determination of vitamin and mineral requirements

To determine vitamin auxotrophy (**Figure S7**), twelve variations of DDM media were prepared. All variations lacked WVS and contained 9 out of 10 individual vitamins, which were prepared as 0.1 mg/ml aqueous stock solutions and added to final concentration equal to that in 5% v/v WVS: (0.1 mg/L (0.41 μM) biotin, 0.1 mg/L (0.23 μM) folic acid, 0.5 mg/L (2.4 μM) pyridoxine-HCl, 0.25 mg/L (0.74 μM) thiamine-HCl, 0.25 mg/L (0.66 μM) riboflavin, 0.25 mg/L (2.0 μM) nicotinamide, 0.25 mg/L (1.0 μM) calcium-D-pantothenate, 5.0 μg/L (3.7 nM) cyanocobalamin, 0.25 mg/L (1.8 μM) p-aminobenzoic acid, 0.25 mg/L (1.2 μM) *α*-lipoic acid). Cells were pre-grown in rPHG, washed and subcultured 1:100 in 50 ml DDM without WVS and grown to stationary phase. Starved cells were washed, diluted and used for growth curves as previously described. To validate requirements for specific vitamins observed during auxanography (**Figure 4**), cells were first starved of vitamins as described and subsequently inoculated into DDM without WVS but containing combinations of biotin, thiamine, cyanocobalamin and L-methionine. We included trace metals (provided by WMS) in our defined media formulations due to the use of highly-purified molecular grade water in our experiments, and also to provide metal-replete conditions inclusive to the potential requirements of yet unknown enzymatic functions, particularly in the context of redox reactions, oxygen tolerance and lignocellulose degradation. WMS can be diluted 100-fold (0.01% v/v WMS) relative to the standard amount in DMM (1% v/v WMS) with no change in growth phenotype in defined media (**Figure 2**). If desired, trace metals may be omitted from media, particularly in the presence of complex components such as yeast extract or if using distilled or impure water sources.

**Figure 4.**
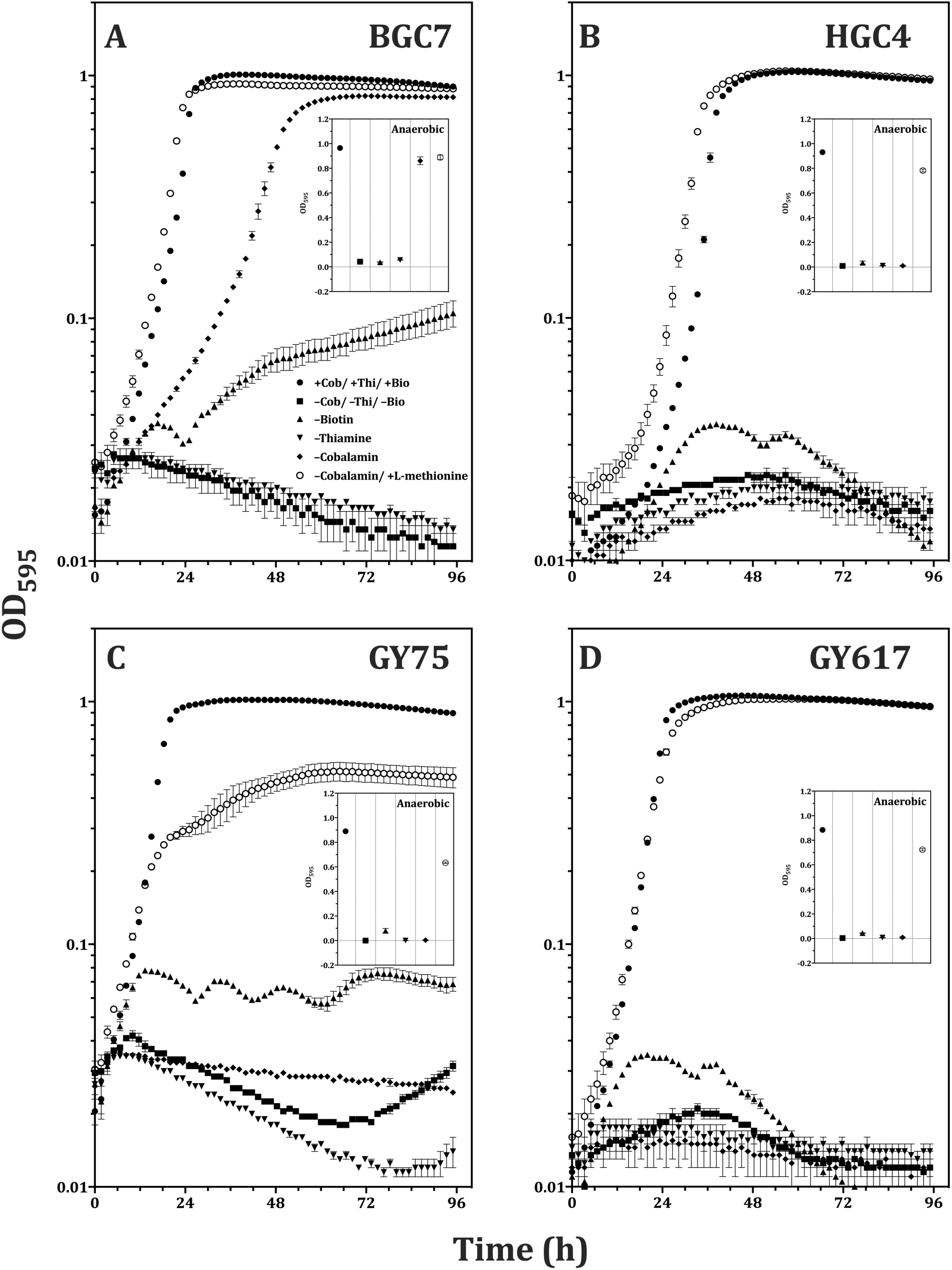
*Dysgonomonas* exhibit growth requirements for thiamine, biotin and cyanocobalamin. *Dysgonomonas* isolates BGC7 (A), HGC4 (B), GY75 (C) or GY617 (D) were pre-grown in DDM lacking vitamins, washed and diluted into 1X M9 salts supplemented with 1% v/v WMS, 0.5% w/v D-glucose, 1.7 mM L-cysteine, 10% v/v hemin solution, 50 μg/ml kanamycin sulfate, pH 7.5, containing 0.41 μM biotin, 0.74 μM thiamine, 3.7 nM cyanocobalamin and 0.67 mM L-methionine and amended as described in the legend in panel A. Insets show final corrected OD_595_ under anaerobic conditions; the y-axis is in linear units. Error bars represent standard error of the mean (n=2).

### Serial culture in DMM

To confirm that DMM contained all nutritional requirements, cultures were serially transferred in DMM (**Figure 5**) as follows. Isolates were pre-grown anaerobically in 5 ml volume of liquid rPHG, washed, diluted as previously described and inoculated 1:20 into 200 μl freshly prepared DMM in 96-well microtiter plates. Cultures were grown aerobically to stationary phase (48 hours) and endpoint OD_595_ readings taken after a two-minute orbital shake. Each transfer event reflected a direct, unwashed 1:10 subculture into fresh DMM media. The transfer from rPHG to DMM represented transfer #0 and serial transfers were repeated 10 additional times. Datapoints represent (OD final) − (OD initial) at 595 nm for each transfer. Linear regression analysis was performed using individual datapoints from all replicates and the best fit curves along with 95% confidence intervals were plotted. The slope of the linear fit is provided, along with the p-values indicating the probability of the slope being significantly non-zero.

**Figure 5.**
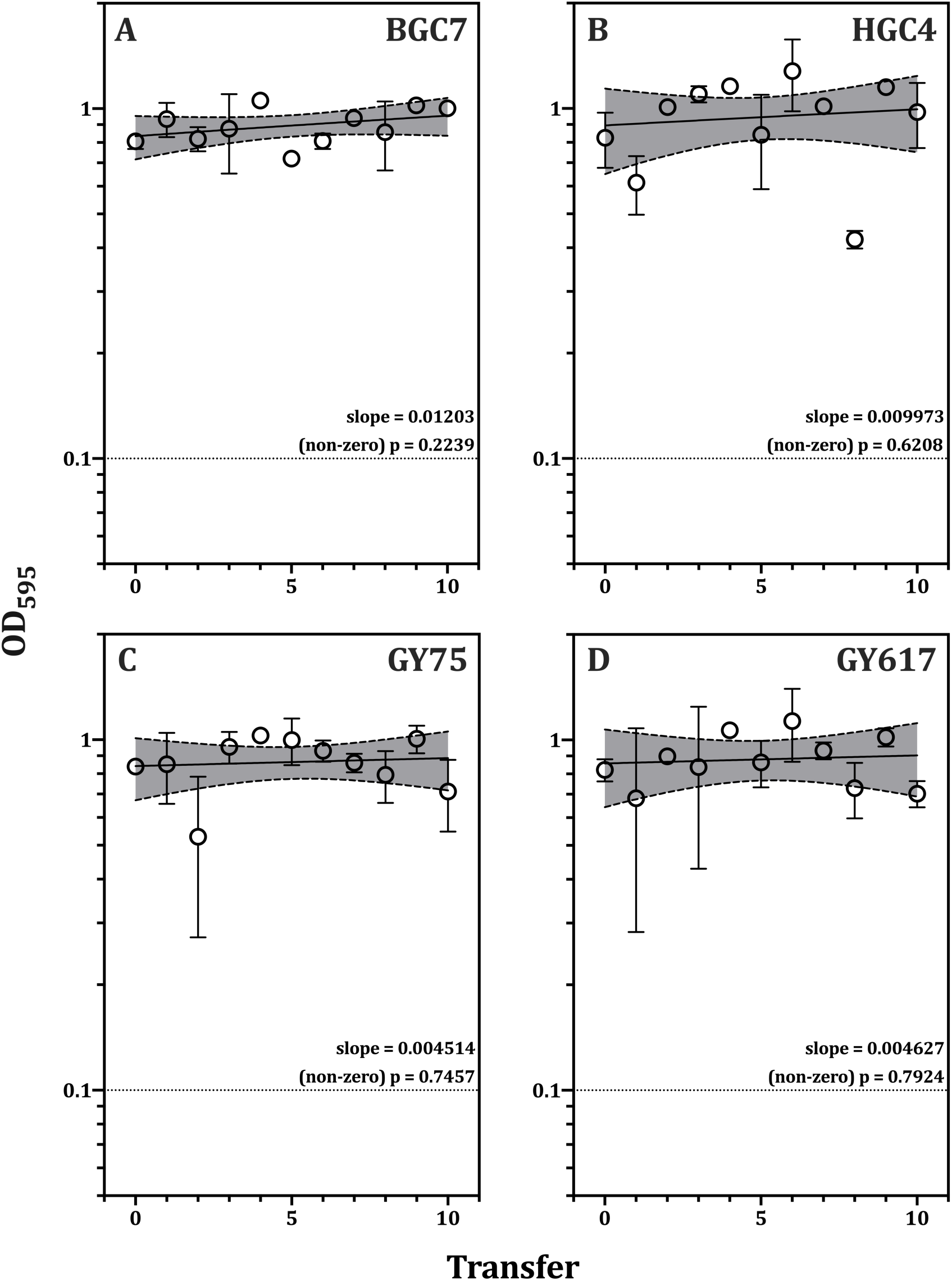
Serial transfer of *Dysgonomonas* in DMM liquid cultures. Cultures were grown to stationary phase in DMM and diluted 1:10 into fresh DMM media and again allowed to reach stationary phase. Cultures underwent a total of 10 serial transfers, where transfer #0 represents inoculation from complex medium. Error bars represent standard error of the mean (n=2). Slopes and their associated p-values (representing significant deviation from zero) for the endpoint readings were generated using linear regression analysis in Prism8.

### Growth kinetics in complex and minimal liquid media

To create a set of interchangeable, parallel liquid media, we used DDM as a basal media to which 10% w/v aqueous solutions of proteose peptone #3 and yeast extract were each added to a final concentration of 1% v/v. To demonstrate expected growth behaviors within and between media, DMM- or DCM-adapted cells were subcultured into either DMM or DCM, as follows. Several DCM-grown colonies of each isolate were used to inoculate 5 ml volumes of liquid DMM and grown to stationary phase as described. DMM cultures were pelleted, washed and subcultured 1:100 into either DCM or DMM and again allowed to reach stationary phase. Subcultures were centrifuged, washed and resuspended as previously described and inoculated separately into DMM and DCM in triplicate. Growth curves were generated as previously described (**Figure 6**). Blank-corrected OD_595_ values for the entire 96 hour duration were used as input data to measure growth kinetics in R [77] package Growthcurver [78] with default settings. Growth rates, generation times and carrying capacities along with standard error values are listed for each isolate in each condition in **Table 1**. Recipes for DMM, DDM and DCM as well as all required components can be found in **Tables S2 & S3**.

**Figure 6.**
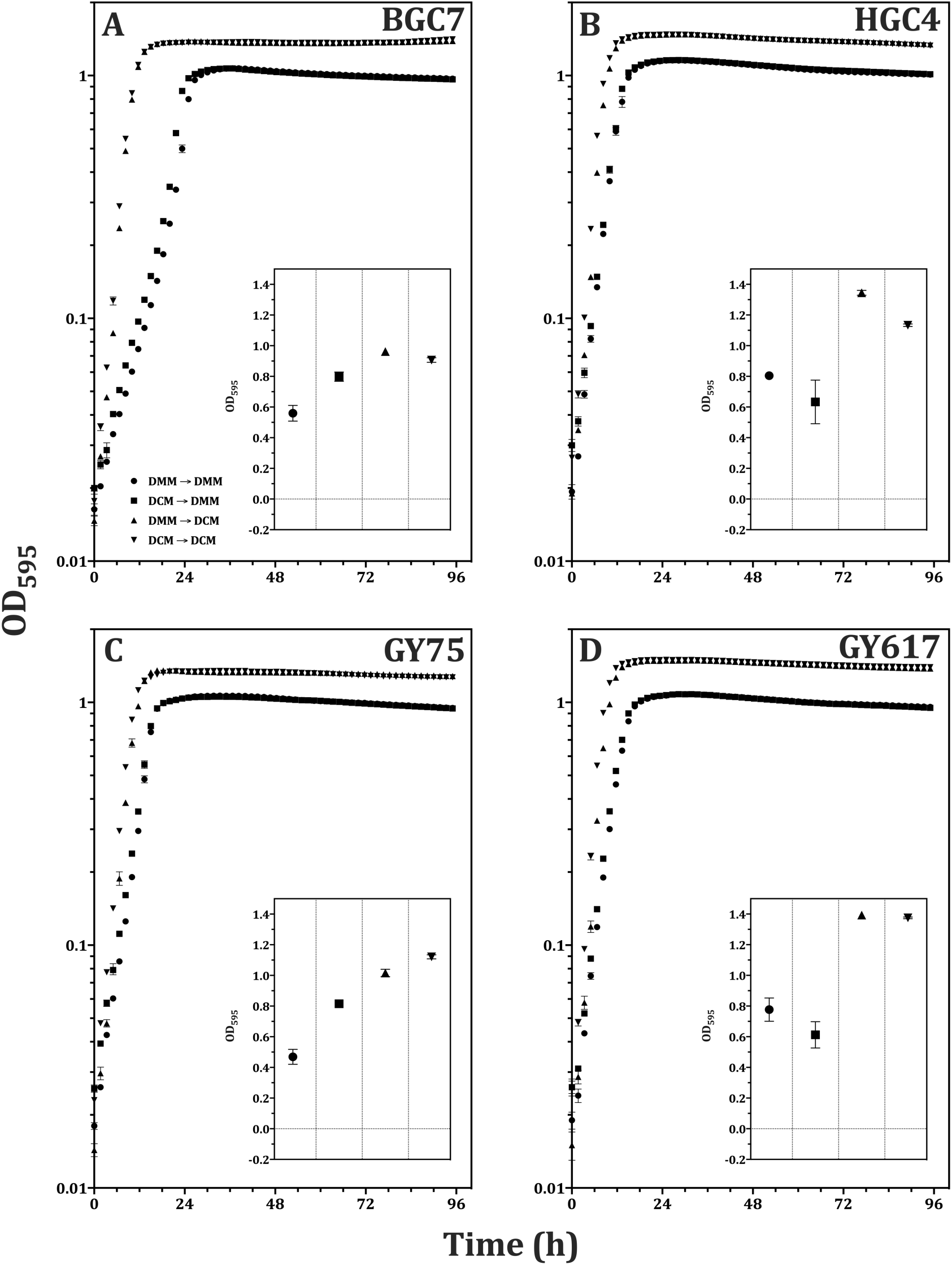
Comparison of growth phenotypes in complex and minimal media. Isolates were pre-grown to stationary phase in either complex (DCM) or minimal (DMM) media and transferred separately into the same or opposite media. Cultures were grown in triplicate to stationary phase aerobically or anaerobically. Pre-growth and growth combinations are located in the legend in panel A. Insets show final corrected OD_595_ under anaerobic conditions; the y-axis is in linear units. Error bars represent standard error of the mean (n=3). Growth rates and carrying capacity were measured using the R package Growthcurver, and results are reported in **Table 1**.

**Table 1.**
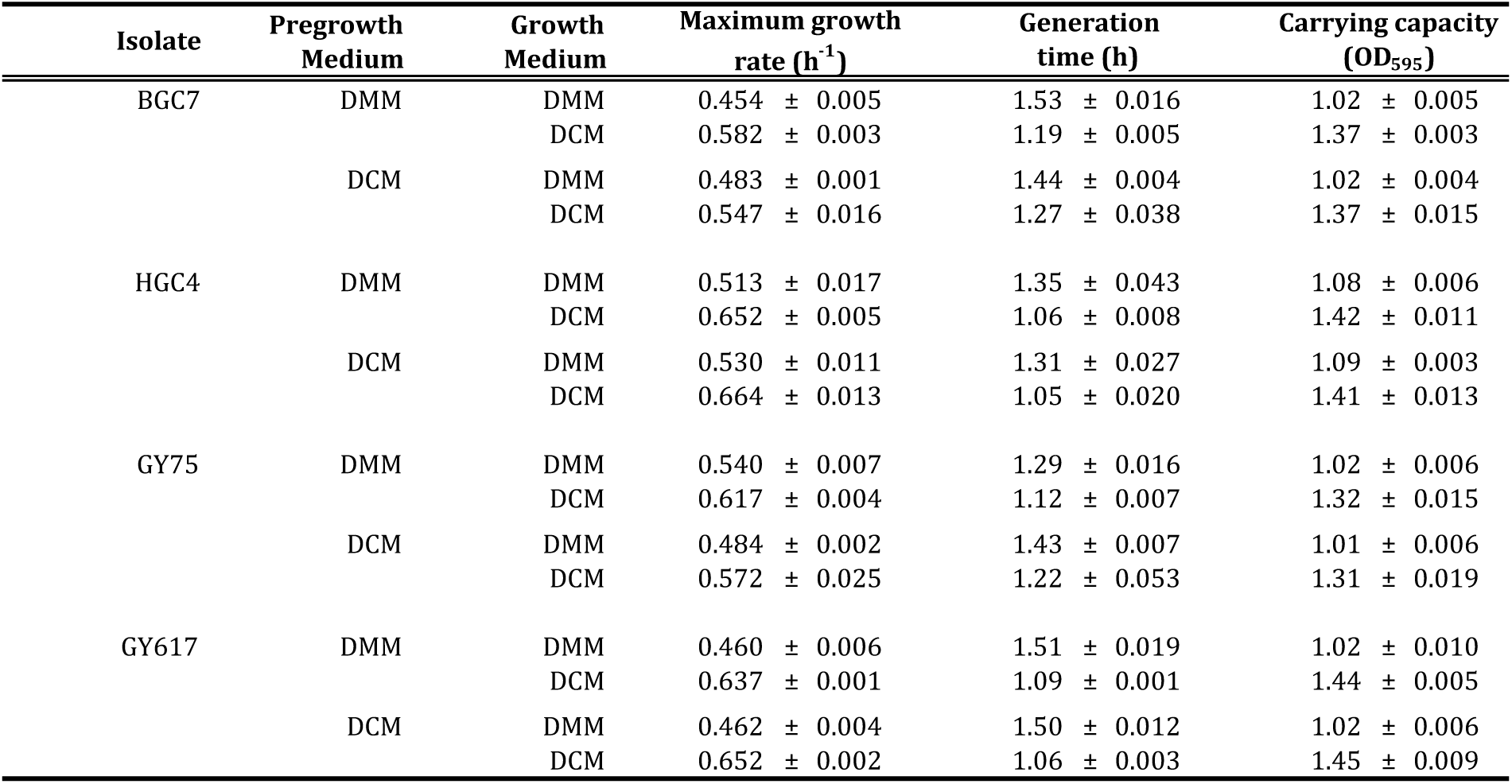
Aerobic growth metrics in DCM and DMM. Isolates were inoculated from solid complex media into DMM broth and allowed to reach stationary phase. Cultures were washed, diluted and subcultured into either DMM or DCM (DMM- or DCM-adjusted cultures) before being washed, diluted and used as inoculum for growth curves (Figure 10). Growth data were fit to the logistic growth model and metrics calculated using the R package Growthcurver. Reported values represent the mean ± standard error (n=3).

| Isolate | Pregrowth Medium | Growth Medium | Maximum growth rate (h<sup>-1</sup>) | Generation time (h) | Carrying capacity (OD<sub>595</sub>) |
| --- | --- | --- | --- | --- | --- |
| BGC7 | DMM | DMM | 0.454 ± 0.005 | 1.53 ± 0.016 | 1.02 ± 0.005 |
|  |  | DCM | 0.582 ± 0.003 | 1.19 ± 0.005 | 1.37 ± 0.003 |
|  | DCM | DMM | 0.483 ± 0.001 | 1.44 ± 0.004 | 1.02 ± 0.004 |
|  |  | DCM | 0.547 ± 0.016 | 1.27 ± 0.038 | 1.37 ± 0.015 |
| HGC4 | DMM | DMM | 0.513 ± 0.017 | 1.35 ± 0.043 | 1.08 ± 0.006 |
|  |  | DCM | 0.652 ± 0.005 | 1.06 ± 0.008 | 1.42 ± 0.011 |
|  | DCM | DMM | 0.530 ± 0.011 | 1.31 ± 0.027 | 1.09 ± 0.003 |
|  |  | DCM | 0.664 ± 0.013 | 1.05 ± 0.020 | 1.41 ± 0.013 |
| GY75 | DMM | DMM | 0.540 ± 0.007 | 1.29 ± 0.016 | 1.02 ± 0.006 |
|  |  | DCM | 0.617 ± 0.004 | 1.12 ± 0.007 | 1.32 ± 0.015 |
|  | DCM | DMM | 0.484 ± 0.002 | 1.43 ± 0.007 | 1.01 ± 0.006 |
|  |  | DCM | 0.572 ± 0.025 | 1.22 ± 0.053 | 1.31 ± 0.019 |
| GY617 | DMM | DMM | 0.460 ± 0.006 | 1.51 ± 0.019 | 1.02 ± 0.010 |
|  |  | DCM | 0.637 ± 0.001 | 1.09 ± 0.001 | 1.44 ± 0.005 |
|  | DCM | DMM | 0.462 ± 0.004 | 1.50 ± 0.012 | 1.02 ± 0.006 |
|  |  | DCM | 0.652 ± 0.002 | 1.06 ± 0.003 | 1.45 ± 0.009 |

### Growth on agar-solidified media

To create parallel, agar-solidified media, single antioxidants were added to agar-solidified DDM according to Dione et al. [79]. Aqueous solutions of reduced glutathione (Fisher, 10 mg/ml) L-ascorbic acid sodium salt (Acros; 10 mg/ml) or uric acid potassium salt (Sigma; 10 mg/ml in 1M NaOH) were prepared and added to 1.5% w/v agar-containing DDM at final concentrations of 0.1 mg/ml, 1 mg/ml or 0.4 mg/ml, respectively. Media was pH-adjusted to 7.5 as necessary using concentrated HCl or NaOH. Isolates were cultured from rPHG-grown glycerol stocks onto agar-solidified complex medium and several colonies used to inoculate liquid DDM cultures. Stationary-phase cells were washed and resuspended to OD 0.1 and spotted onto agar-solidified DDM plates with and without antioxidant amendment. Plates were loosely bagged in plastic petri dish bags (Fisher) and incubated for 12 days aerobically at 22°C and photographed at 7 and 12 days post inoculation (dpi) (**Figure 7**). To test growth on solid DCM, single colonies of each isolate were taken from rPHG plates and streaked to isolation on DCM amended with 1.5% w/v agar and 0.1 mg/ml reduced glutathione. Plates were incubated loosely bagged aerobically at 22°C or 30°C, or anaerobically at 22°C for 4 days (**Figure S8**). To test solid DMM, several colonies of each isolate were taken from solid DDM plates and individually resuspended into 200 μl 1X M9 salts, spotted onto agar-solidified DMM containing 0.1 mg/ml reduced glutathione and streaked to isolation. Plates were incubated with either no bag (unrestricted atmosphere) or sealed in a plastic bag (restricted atmosphere) and incubated aerobically at 30°C for 5 days (**Figure S9**).

**Figure 7.**
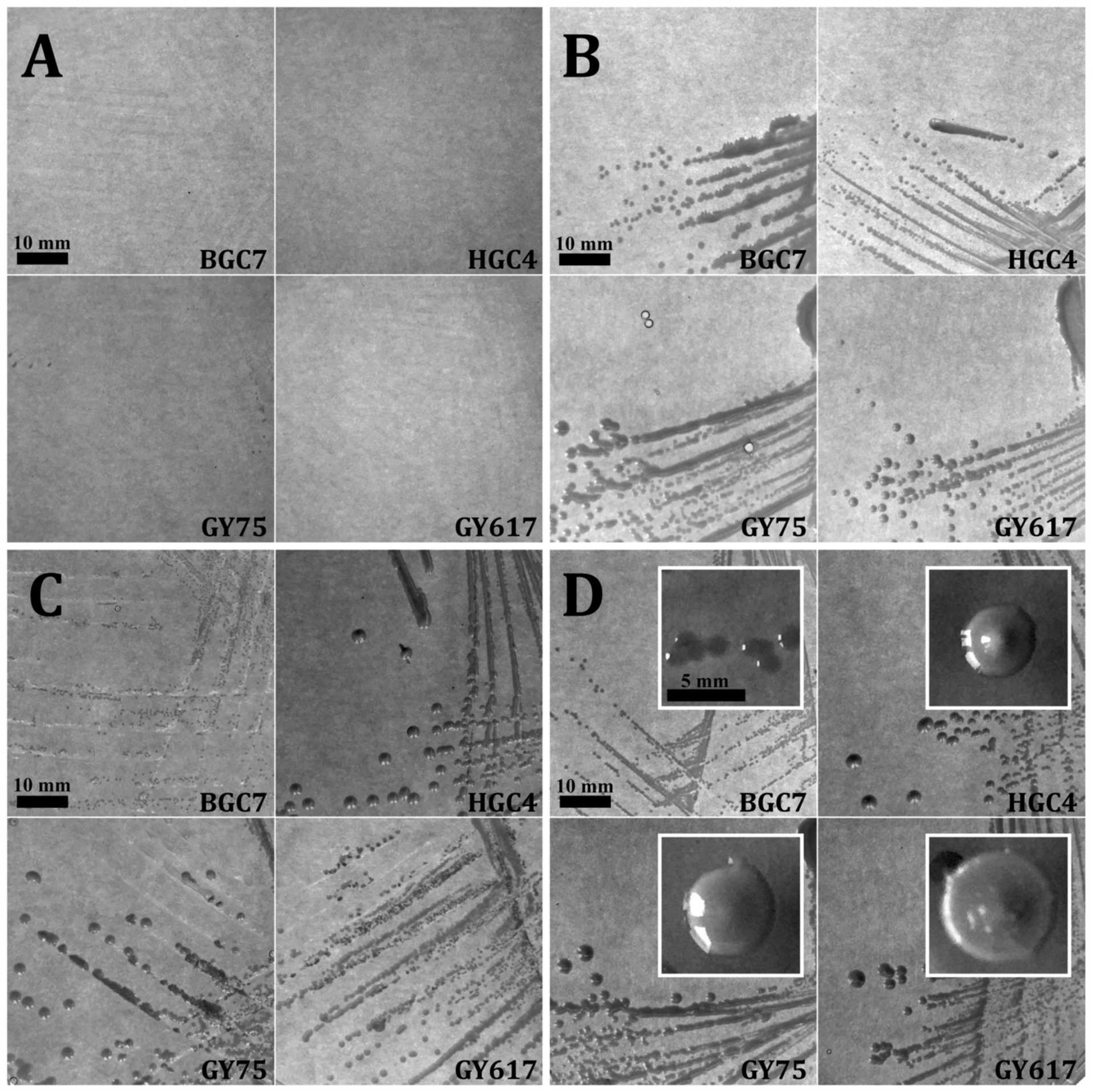
Antioxidants are required for aerobic growth on defined medium. *Dysgonomonas* isolates BGC7, HGC4, GY75 and GY617 were grown on agar-solidified DDM containing: No supplemental antioxidants (A), 0.1 mg/ml glutathione (B), 1 mg/ml L-ascorbic acid (C), 0.32 mg/ml uric acid (D). Overnight cultures were grown in DCM to stationary phase and washed and diluted to OD_595_ of 0.1 and 20 μl spots were allowed to dry onto media before streaking to single colonies. Plates were loosely bagged and incubated aerobically at 22°C for 12 days. Photographs were taken at 7 dpi (main panels) or 12 dpi (insets). Scale bars represent 10 mm (main panels) or 5 mm (insets).

### Enrichment of *Dysgonomonas isolates* from termite hindgut

Termites were collected from the same site and colony as isolates GY75 and GY617 in June 2019 during an alate (winged reproductive) swarming event. Hindguts from freshly collected worker (n=25) and alate (n=50) castes were extirpated and the contents separately prepared and diluted as previously described. Dilutions were plated on agar-solidified DMM containing 0.1 mg/ml reduced glutathione, 50 μg/ml kanamycin sulfate and 100 μg/ml cycloheximide (Acros), loosely bagged and incubated aerobically or anaerobically at 22°C for 4 days. Well-isolated colonies that exhibited morphology characteristic of *Dysgonomonas* (circular, convex, opaque, glossy, white-cream or light-brown in color) were purified by serially streaking onto the same medium, and 30 isolates were selected for identification. We used a high-throughput screen to quickly identify the presence of *Dysgonomonas* spp. within the collection of isolates. Crude gDNA was prepared individually for each isolate, in which several colonies were resuspended in 100 μl sterile molecular-grade water, boiled at 95°C for 15 minutes and briefly centrifuged. The supernatant was used as template for PCR targeting the V4 region of the bacterial 16S rRNA gene. Each isolate was individually PCR amplified using Q5 Polymerase and primers 16S_F (5’-GTGCCAGCMGCCGCGGTAA-3’) and 16S_R (5’-GGACTACHVGGGTWTCTAAT-3’), each flanked by 5’ extensions containing unique combinations of forward or reverse indexing sequence and high-throughput sequencing adapters [80]. The negative control replaced lysate supernatant with water. Positive controls included lysate supernatant from *Dysgonomonas* spp. BGC7 & HGC4 and isolates GY75 & GY617. Amplicons were pooled, column-purified and sent to the UConn MARS Facility (University of Connecticut, Storrs, CT, USA) where libraries were prepared and sequenced on the Illumina MiSeq platform. Raw reads were processed in R using the package DADA2 [81] to quality filter and trim, merge, chimera-check, determine read error rates, generate Amplicon Sequence Variants (ASVs) [82] and to assign taxonomy to ASVs using the SILVA rRNA database v132 [83]. Taxonomic assignments were manually inspected and preliminary identification was determined by the taxonomic group with which >95% of reads were placed, with the remaining reads attributed to PCR and sequencing errors that passed the filtering process. Isolates that were identified as belonging to genus *Dysgonomonas* (24 of 30), along with the controls, clustered into 4 discrete ‘ASV-groups’ (ASV_1, ASV_2, ASV_4, and ASV_5), each of which contain 253 bp sequences with 100% identity with each other. ASVs from this study can be found in **Table S4**. To aid in resolving subtle variations in 16S rRNA gene heterogeneity within ASV-groups, a subset of the isolates (13 of 24) containing representatives from each ASV-group were selected for bidirectional dideoxy sequencing to obtain full length 16S rRNA gene sequences. gDNA template was prepared and quantified for each isolate within the subset. PCR, cloning and sequencing were carried out as previously described. Sequences were trimmed of primer sequence and base-calls manually inspected and curated using the generated ASV sequences as a reference. Full-length 16S rRNA gene sequences for *Dysgonomonas* isolates obtained or used in this work (AAn1, AAn3, AAn4, AAn6, AAn7, AAn9, AAn11, BGC7, GY75, GY617, HGC4, WAe4, WAe5, WAe6, WAe3, WAn2, WAn3) were submitted to GenBank under accession numbers MT340871-MT340887, respectively. Isolates were named by isolation source (<u>W</u>orker or <u>A</u>late), oxygen condition (<u>Ae</u>robic or <u>An</u>aerobic), and isolate number. For example, isolate AAn1 was isolated from alate hindguts under anaerobic conditions, while WAe3 was isolated from a worker hindgut under aerobic conditions. Colonies with morphology consistent with that of *Dysgonomonas* were obtained aerobically from alate hindguts (A<u>Ae</u>) but were discarded due to fungal overgrowth. Reference sequences from cultured members of *Dysgonomonas* and from representative taxa from within the *Bacteroidetes* greater than 1.2 kb in size were obtained from RefSeq or GenBank, in order of preference, on April 8, 2020. Reference sequences were trimmed of primers and aligned with those from *Dysgonomonas* spp. BGC7 and HGC4 and isolates from this study using the MUSCLE algorithm in Geneious R9 with default settings (see **File S1** for multiple sequence alignment of 16S genes and **File S2** for pairwise nucleotide identifies). IQ-TREE v2.0 [84] was used for substitution model testing and maximum likelihood phylogenetic reconstruction using model TIM3e+I+G4. Branch support values were calculated using 1000 Ultrafast Bootstraps and SH-aLRT testing. Isolation sources for *Dysgonomonas* isolates not from this study were obtained from the published reference when available or from data associated with the RefSeq/GenBank entry. Formatting and metadata were applied in R using packages dplyr [85], ggplot2 [86], treeio [87], ggtree [88], and ggrepel [89]. Explanations of formatting and metadata can be found in the legend for **Figure 8**.

**Figure 8.**
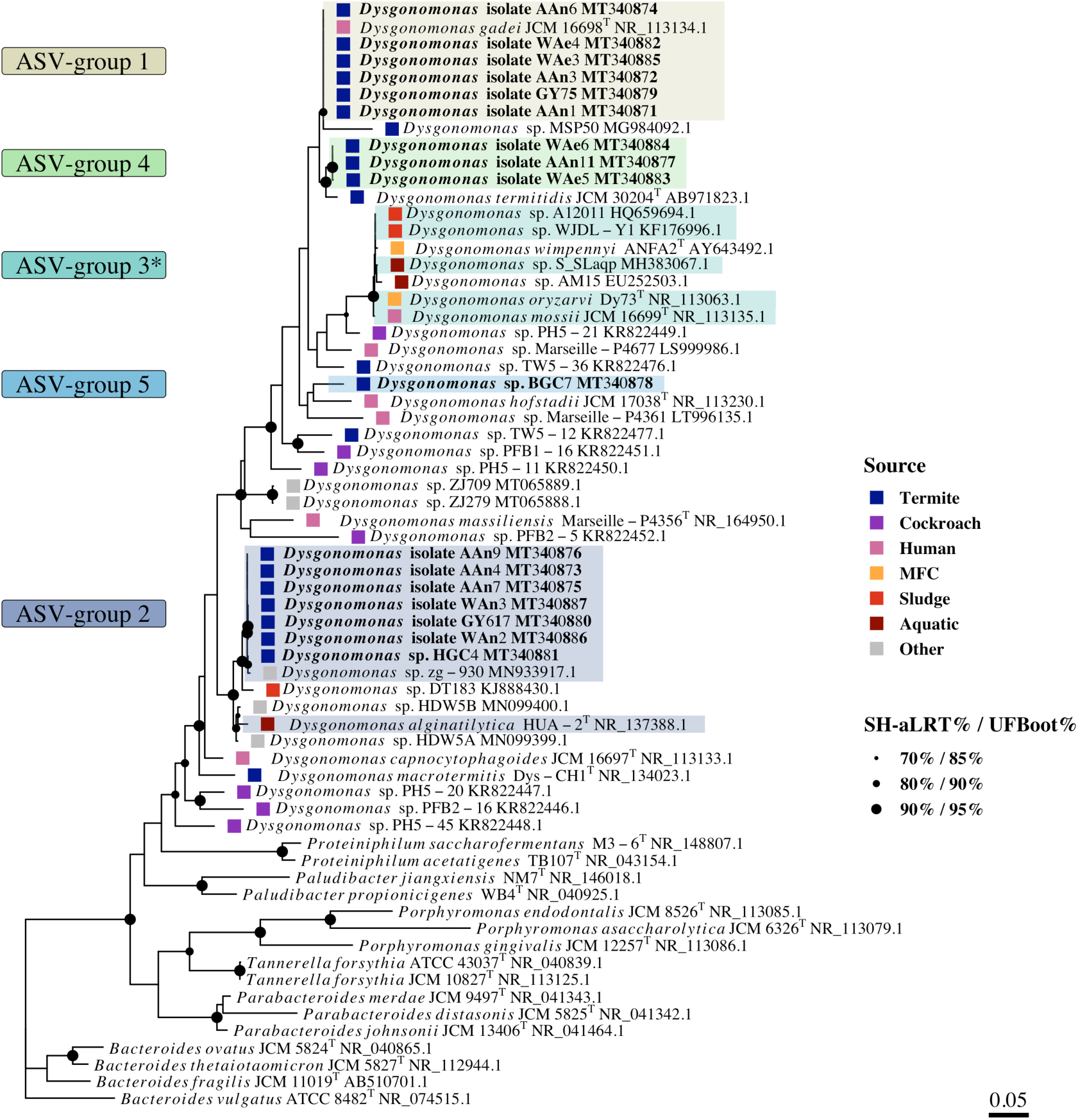
Maximum-likelihood phylogenetic reconstruction of near-full length 16S rRNA gene sequences from cultured members of genus *Dysgonomonas*. Sequences of length 1.2-1.5 kb were acquired from RefSeq and GenBank and combined with quality-controlled sequences from *Dysgonomonas* isolates from this study. Nodes containing filled circles represent branch support values; small, medium or large node shapes represent 70%/85%, 80%/90% or 90%/95% SH-aLRT/UFBoot support, respectively. The scale bar represents 0.05 substitutions per site. Type strains are designated by a superscript ‘T’ following the strain designation. Isolates used or obtained in this study are bolded. Bolded *Dysgonomonas* isolates beginning with ‘A’ are derived from alates while those beginning with ‘W’ are worker-derived. Highlighted clades represent ASV-groups, which share common ASVs (contain 100% identity over the 250 bp sequence containing the V4 region of the 16S rRNA gene). *Dysgonomonas* spp. not obtained or used in this study were included as part of an ASV-group if they also exhibited 100% identity to ASVs generated in this study. ASV-group 3* represents taxa which share 100% identity over the same 250 bp sequence of 16S rRNA gene, but were not true ASVs generated in this study. Square tip points are colored by isolation source according to the legend. Source category ‘Other’ contains *Dysgonomonas* isolates from marmot (MT065889.1), bird (MT065888.1), crayfish (MN933917.1) and beetle (MN099400.1 & MN099399.1).

## Results and discussion

### Growth in complex vs. defined media

We tested four phylogenetically diverse termite-derived *Dysgonomonas* isolates for their ability to grow in aerobic, liquid media consisting of M9 salts supplemented with trace minerals, ferric hemin, B-vitamins, D-glucose and proteose peptone (**Figure 1**). Previous experiments determined that removal of peptone, and thus amino acids, from the medium resulted in no growth from all isolates (data not shown), suggesting a growth requirement for one or more components found in peptone. To determine which components of peptone were necessary to sustain growth, cultures were supplied with B-vitamins (WVS) and trace metals (WMS), and proteose peptone was replaced with either casamino acids or a mixture of twenty individual free amino acids, each at 0.1 mg/ml. Casamino acids were supplemented with 0.1 mg/ml L-tryptophan and L-cysteine to replenish amino acids lost during acid hydrolysis. In comparison to growth in a medium with peptone, isolates BGC7, HGC4 and GY617 exhibited slight decreases in growth rate and yield when grown with casamino acids. Isolate GY75 displayed growth kinetics most comparable to that on peptone, demonstrating a very slight decrease in growth rate, and no discernible reduction in yield. Free amino acids were able to be substituted for peptone or casamino acids for all isolates, though growth rates and yields were decreased. Additionally, isolates HGC4 and GY617 exhibited a slight diauxie-like phenotype when grown with casamino acids which was exacerbated by growth on free amino acids. This phenotype may be attributed to differences the relative abundance of particular amino acids or oligopeptides between the supplements. Nucleobases, NAD and non-proteinogenic amino acids L-ornithine and DAP were not required for growth, nor did they contribute to additional yield increases beyond that of free proteinogenic amino acids alone. Nucleobases had a negative effect on all measures of growth from isolate BGC7, perhaps due to interference between biosynthesis and salvage pathways. Together these results suggested that growth could occur on oligopeptides or free amino acids in the presence of trace metals and B-vitamins, and suggested a possible amino acid auxotrophy.

### Amino acid auxanography

Amino acid auxanography was performed to determine the growth requirement for one or more amino acids. Isolates HGC4, GY75 and GY617 exhibited growth only on amino acid pool 2 (L-alanine, L-cysteine, L-threonine, L-asparagine, L-methionine, DAP) & pool 6 (L-serine, L-cysteine, L-ornithine, glycine), which suggested L-cysteine as the single limiting amino acid (**Figures S1 & S2**). Additionally, isolates HGC4 & GY617 again exhibited a diauxie-like growth phenotype, likely indicating limitation of one of more preferred nutrients in the auxanography growth medium. Isolate BGC7 exhibited growth on pools 2, 6 & 9 (L-methionine, L-glutamic acid, L-lysine), with greatest yield on the L-cysteine- and L-methionine-containing pool 2. Pool 6, which contained L-cysteine but not L-methionine, was growth-permitting for isolate BGC7, but both growth rate and yield suffered. Isolate BGC7 also exhibited weak, linear growth on pool 9, suggesting that L-methionine alone, while not optimal, is growth-permitting under the conditions tested (**Figure S2**). These results suggest that L-cysteine alone or in combination with L-methionine can replace complex sources of amino acids and allow exponential aerobic growth of *Dysgonomonas*.

### L-cysteine is required during growth under aerobic or anaerobic conditions

To examine relationship between growth kinetics and L-cysteine, isolates were grown aerobically and anaerobically across a range of L-cysteine concentrations. Freshly prepared trace metals, B-vitamins and ferric hemin were provided and L-cysteine supplemented between 0-5.69 mM. All cultures exhibited concentration-dependent phenotypes in relation to L-cysteine (**Figure S3**) under both aerobic and anaerobic conditions. With respect to growth rate under aerobic conditions, isolates displayed either a positive correlation with L-cysteine concentration (isolate BGC7) or a binary response (isolates HGC4, GY75 & GY617) in which nearly maximal growth rate was achieved at the lowest concentration tested (0.285 mM). When L-cysteine was absent from aerobic media, isolates exhibited extended lag phase, periods of linear growth and early entry to stationary phase suggestive of L-cysteine limitation. Similar to the results from amino acid auxanography, isolate BGC7 grew modestly under aerobic conditions in the absence of L-cysteine and exhibited extended lag phase, low growth rate and low yield. Isolate BGC7 exhibited reduced lag time and an increase in growth yield, as well as an increase in growth rate that was positively correlated with L-cysteine concentration up to 3.42 mM, beyond which no gains in growth rate, or yield, were obtained. Anaerobic growth of isolate BGC7 showed positive correlation between L-cysteine concentration and final yield up to 4.56 mM. Aerobically, isolate HGC4 and isolates GY75 & GY617 did not exhibit the same relationship between L-cysteine and growth rate as isolate BGC7, and instead displayed a binary response which required only 0.285 mM to achieve near-maximal growth rates and yields. For isolates HGC4 and GY617, 0.285 mM L-cysteine was yield-limiting, but 0.596 mM was considered L-cysteine replete under the conditions tested. Anaerobic growth was similarly binary for these isolates, with no growth occurring in the absence of L-cysteine. The replacement of peptones or casamino acids with L-cysteine as a component of the basal salt-vitamin-hemin-glucose medium led to the development of a chemically defined medium, which was later refined to become DDM.

As previous experiments demonstrated L-cysteine-dependent growth, we sought to determine whether pre-growth in rich medium provided the means for cells to intracellularly store sulfur and permit moderate growth in media without L-cysteine. Forthcoming draft genome assemblies for *Dysgonomonas* isolates BGC7, HGC4, GY75 & GY617 all contain genes encoding sulfate permease, but pathways for assimilatory sulfate reduction are absent. As such, we were further interested to determine whether sulfate was necessary in the presence of a reduced, assimilable source of organic sulfur such as L-cysteine. Cultures were pre-grown to stationary phase in DDM with 0.57 mM L-cysteine, washed several times and diluted into PBS. Prepared cells were diluted 1:100 into fresh media with and without sulfate or L-cysteine and grown under aerobic and anaerobic conditions. Sulfate-limited DDM (0.01% v/v mWMS) contained 83 nM total sulfate, which is ∼300-fold less than the required 26.5 μM for *Salmonella typhimurium* to achieve an OD_420_ of 1 in MOPS medium [90], while sulfate-replete DDM (0.01% v/v WMS) contained 1 mM total sulfate. L-cysteine concentrations were either 0 or 1.71 mM, and kanamycin sulfate was omitted from all media. It should be noted that both sulfate-replete and sulfate-limited DDM contain the vitamins biotin, thiamine and lipoic acid (found in WVS). These cofactors contain organic reduced sulfur, and together contribute ∼1.15 μM to the pool of putatively available reduced sulfur. All isolates failed to grow either aerobically or anaerobically in the in the absence of L-cysteine, regardless of the presence of sulfate in the medium, which suggested a requirement for a reduced form of assimilable sulfur (**Figure 2**). These results further suggested that the growth of isolate BGC7 in the absence of L-cysteine observed in **Figures S2 & S3** may be attributed to intracellular sulfur stores present as a consequence of pre-growth in complex medium. Aerobic growth phenotypes were nearly identical for all isolates in the presence of L-cysteine, regardless of the presence of sulfate in the medium, which suggested that sulfate is not required aerobically in the presence of L-cysteine. Although sulfate was unable to replace L-cysteine as a sulfur source, anaerobic growth was stimulated by the presence of sulfate, particularly for isolate HGC4 and isolate GY75. The role of sulfate during anaerobic growth is unclear, but it is perhaps used as an alternative electron acceptor during anaerobic respiration. All isolates were able to utilize the tripeptide glutathione (glutamate-cysteine-glycine) to some degree in place of L-cysteine, and isolate BGC7 was further able to utilize L-methionine as a sulfur source. No growth occurred when L-cysteine was replaced with thiosulfate, thioglycolate, 2-mercaptoethanol or dithiothreitol (not shown). Sodium sulfide was not tested, but was found to be a suitable sulfur source for *Bacteroides fragilis* [67].

The precise roles of L-cysteine are difficult to discern over the range of concentrations tested here (0.1-10 mM), as it can be imported and used directly as a substrate for peptide synthesis, as a source of assimilable reduced sulfur [67], or as a reducing agent [79, 91] to lower the redox potential of the medium. Draft genome assemblies contain complete L-cysteine and other proteinogenic amino acid biosynthesis pathways suggesting that the isolates are not amino acid auxotrophs, and the ability of L-cysteine concentration to modulate final growth yield also suggests that it is an exhaustible nutrient, assimilated into biomass, or both. Moreover, the requirement for L-cysteine anaerobically suggests that its function as a reductant provides a minor contribution to its role in growth. The inability of other sulfur-containing reducing agents to permit growth may be due simply to the inability of *Dysgonomonas* to effectively assimilate reduced sulfur under the provided conditions. Taken together, these results suggest that L-cysteine is utilized as an easily assimilable source of reduced sulfur, and that other roles that it may provide are secondary to this function.

### Growth using L-cysteine as the sole source of nitrogen

To determine if L-cysteine could also be utilized as a sole source of nitrogen, cultures were grown in the presence or absence of excess ammonium and/or 1.7 mM L-cysteine, which should not be significantly yield-limiting as the sole source of nitrogen [90] under these conditions. All isolates were able to grow in the absence of ammonium, using L-cysteine as a sole source of nitrogen under aerobic and anaerobic conditions (**Figure S4**). Using L-cysteine as a nitrogen source, isolate GY75 exhibited a 24-hour lag before exponential growth using L-cysteine under aerobic conditions. Isolates BGC7, HGC4 & GY617 exhibited similar growth rates to their counterparts grown with ammonium and L-cysteine, but entered stationary phase early, perhaps due to nitrogen limitation or buildup of toxic end products of L-cysteine catabolism such as hydrogen sulfide [92]. Under aerobic conditions in the absence of both ammonium and L-cysteine, some weak growth occurred aerobically, but not anaerobically, which could be indicative of scavenged and assimilated trace nitrogen from the medium (hemin, 0.3 mM; kanamycin, 0.34 mM; WMS, 0.078 mM; biotin, 0.82 μM; thiamine, 2.96 μM; cyanocobalamin, 51.6 nM; totaling ∼0.7 mM nitrogen) or that the growth was enabled by utilizing intracellular nitrogen stores. Further analysis of nitrogen sources was precluded by the requirement for L-cysteine for growth and its ability to be used as a nitrogen source. Under anaerobic conditions, isolates exhibited approximately half the yield as compared to those grown under ammonium-replete conditions. We speculate that assimilation of organic nitrogen could be unfavorable during anaerobic growth under the provided conditions.

### Requirement for ferric hemin

Our *Dysgonomonas* isolates had previously been observed to require ferric hemin when grown under aerobic or anaerobic conditions, even when grown in complex media such as rPHG (author observation). Hemin, a ferric-iron carrying porphyrin, is biosynthesized through a costly and complex pathway, making it a valuable commodity. Respiration requires both sufficient quantities of iron for redox reactions, and iron-containing porphyrins for synthesis b-type cytochromes, [67, 90, 93] which hemin could provide. We sought to determine growth-limiting concentrations of hemin for our *Dysgonomonas* isolates when D-glucose was used as a carbon source (**Figure S5**). Cultures were hemin-starved and prepared as described in Materials & Methods before being used for growth curves. Hemin was present in media at final concentrations between 0 and 153.4 μM (0-20% v/v hemin solution). Similar to that observed in *Bacteroides fragilis* [94], growth of *Dysgonomonas* under aerobic conditions exhibited a positive correlation between hemin concentration and growth yield, up to the growth-saturating concentration for that organism where hemin became excessive and yield remained constant. Isolates did not grow when hemin concentration was below 7.67 μM (1% v/v hemin solution), demonstrating a hemin requirement, though growth-permissive concentrations differed among the isolates tested. All isolates also exhibited poor growth at their respective lowest growth-permissive concentration. Poor growth included extended lag time, slower growth rate and lower yield. Isolates BGC7 and HGC4 required 38.4 μM hemin for growth to occur, with 61.3 μM sufficient for near-maximal growth rate and yield. Isolate GY75 required the least ferric hemin for growth (7.67 μM) and again achieved near-maximum growth rate and yield in the presence of 38.4 μM. Isolate GY617 required the greatest concentration of ferric hemin (46.0 μM) for growth to occur, and near-maximal growth rate required at least 76.7 μM hemin. Ferric hemin had no deleterious effects on growth of any isolates at the higher concentrations tested under these conditions. Hemin requirements may change depending on carbon source, particularly if growth requires extensive oxidation of substrates performed by iron-cofactor dependent electron carriers.

Under anaerobic conditions, the hemin concentration-yield relationship was consistent with a third-order polynomial in which concentration was positively correlated with yield at lower concentrations, but yields either plateaued or decreased over a range of concentrations before reaching maximal yield in the presence of 153 μM hemin (**Figure S6**). Though all isolates exhibited the greatest yield in the presence of 153 μM hemin (20% v/v hemin solution), isolates BGC7, HGC4, GY75 and GY617 each also exhibited a secondary, local maximal yields at 46.0 μM, 30.67 μM, 46.0 μM and 76.7 μM hemin, respectively. Lower concentrations of hemin were required under anaerobic than under aerobic conditions for all isolates, consistent with a switch from aerobic respiration to fermentation, which could require fewer iron-containing electron carriers and respiratory cytochromes. Isolates HGC4 and GY617 required 15.4 μM to exhibit growth, roughly half of that required aerobically. Isolate BGC7 required only 3.83 μM hemin but growth was limited to ∼2.5 generations. Isolate GY75 was able to grow in the absence of hemin, consistent with having the least requirement for hemin during aerobic growth, but was growth was limited to under two generations. It is noteworthy that all isolates can exhibit characteristics consistent with hemin accumulation during growth in hemin-replete media similar to that of close relative *Porphyromonas gingivalis* ([95] and references therein), such as the formation of brown cell pellets from liquid culture or darkening of colonies on agar-solidified media. Isolate GY75 routinely exhibits the greatest cell darkening (author observation) and its ability to grow using the least amount of hemin of the four isolates under aerobic and anaerobic conditions is consistent with the ability to sequester ferric hemin within the periplasm or at the outer cell membrane.

We further sought to determine whether ferric hemin was able to be replaced with alternate sources of iron (**Figure 3**). In addition to ferric hemin, we tested soluble ferric chloride and two ferric chelates; ferric-EDTA and ferric-citrate, as well as soluble ferrous sulfate alone. All conditions were considered replete for ferrous iron by the addition of 1% v/v WMS which provided 3.6 μM ferrous sulfate, the standard for DDM. For all isolates, ferrous iron alone was insufficient to permit growth, and similar to *Prevotella intermedia* [96], none of the soluble or ferric chelates were able to replace ferric hemin under aerobic or anaerobic conditions. Hemin was ultimately used at 76.7 μM (10% v/v hemin solution) in our final media recipes as it was not significantly rate- or yield-limiting and reduced the risk for heme-toxicity which can occur at high concentrations [93]. Although beyond the scope of this study, it would be interesting to determine the ability for *Dysgonomonas* utilize non-heme iron in the presence of alternate porphyrins such as protoporphyrin IX such as related bacteria such as *P. intermedia* [96] *P. ruminicola* [97], *P. gingivalis* [98] and are able.

### Vitamin requirements

Previous experiments determined that WVS was required for growth by all isolates when grown without peptone (data not shown), indicative of auxotrophy for one or more B-vitamins. To screen for vitamin auxotrophies, cultures were vitamin-starved and prepared as previously described before being inoculated into 12 different B-vitamin-replete media, each with an omission of a single cofactor (**Figure S7**). Removal of biotin, thiamine or cyanocobalamin limited growth from all isolates with the exception of isolate BGC7, which exhibited significantly slower growth rate but similar yield in the absence of cyanocobalamin. All isolates were able to grow to some degree in the absence of biotin, which suggested intracellular reserves in the inocula were not yet biotin-limited during the vitamin pre-starvation procedure. All isolates exhausted their intracellular biotin stores through cell growth and dilution within a few generations and growth ceased. We subsequently tested the requirements for these three cofactors by omitting WVS from the medium, amending only with biotin, thiamine and cyanocobalamin and then removing each cofactor individually (**Figure 4**). Consistent with previous findings, all isolates displayed the expected growth phenotypes when provided biotin, thiamine and cyanocobalamin together. Omission of all vitamins resulted in absence of growth from all isolates. Thiamine and biotin were required for the growth of all isolates, while cyanocobalamin was required for all but isolate BGC7. Some growth was observed from all isolates in the absence of biotin, again suggesting that cells in the vitamin-starved inocula contained an exhaustible intracellular pool of biotin. Isolates also exhibited some degree of oscillatory growth in the absence of biotin alone. For example, isolates HGC4 and GY617 ultimately entered a death phase after oscillation, while isolate BGC7 continued a pattern of linear growth, indicative of cofactor limitation. Isolate GY75 exhibited oscillatory growth in the absence of biotin for the duration of the 96-hour assay. Cyanocobalamin was required for growth for all strains except BGC7, which exhibited a lower growth rate but nearly equivalent yield compared to cells grown with cobalamin. Cobalamin-independent growth of isolate BGC7 was confirmed by diluting washed cells 1:100 into fresh medium without cyanocobalamin and allowed to reach stationary phase. Cells were able to be serially passaged three times in this manner (not shown), which confirmed that isolate BGC7 can indeed grow in the absence of cyanocobalamin. Genomic data show the absence of de novo cobalamin biosynthetic pathways, but we cannot rule out the ability of isolate BGC7 to biosynthesize cobalamin through non-canonical pathways. It also cannot be ignored that hemin and cobalamin share the intermediate compound uroporphyrinogen-III, which could be, although energetically costly, created by dissimilation of hemin. This conundrum has been previously explored in *Porphyromonas gingivalis* [98], although clear answers were also not derived. Alternatively, it is possible that isolate BGC7 could grow in the absence of cobalamin due to the presence of genomic features such as cobalamin-independent methionine synthase, for example. In the presence of thiamine and biotin, the substitution of cyanocobalamin with L-methionine allowed high growth rates for all isolates, although the provided 0.67 mM L-methionine appeared yield-limiting for isolate GY75. This result suggests that although cyanocobalamin may be required to perform multiple functions, a major role for cellular cobalamin during aerobic and anaerobic growth of some *Dysgonomonas* species is likely biosynthesis of L-methionine [99].

### Serial culture in DMM

To be certain that DMM did not lack growth factors required only in trace amounts, we performed ten serial aerobic transfers of four *Dysgonomonas* isolates in DMM. Cells from previous DMM cultures were diluted 1:10 into fresh DMM without washing and allowed to reach stationary phase before being transferred again. Each transfer allowed contaminating nutrients that were either intracellular, bound to the cell surface or carried along with culture supernatant to be diluted into fresh medium, and then diluted by redistribution amongst actively growing cells within the population. All four isolates were able to be serially transferred ten times in DMM without a loss in final yield (**Figure 5**). Cultures were not shaken during these experiments and thus endpoint OD_595_ readings did exhibit some fluctuation due to cell clumping and biofilm formation. Linear regression analysis demonstrated that the slope of each fit to final OD_595_ values was positive, which indicated that final yield did not decrease during successive transfers. The linear fit was tested for significant deviation from zero, and all isolates exhibited p-values >0.22, which indicated that the slopes of the linear fits were not significantly different from zero. Taken together these data confirmed that liquid DMM was indeed sufficient to meet the growth needs for the *Dysgonomonas* isolates used here. Although we tested *Dysgonomonas* isolates with diverse phylogenetic placement, it is possible that other strains may exhibit auxotrophy for specific amino acids or vitamins. Using DMM as a basal minimal medium to perform further amino acid or cofactor auxanography will allow simple and quick elucidation of any alternate growth requirements.

### Growth kinetics in DCM & DMM

To create a parallel complex media, we amended DDM (B-vitamin replete) with 1% v/v each of peptone and yeast extract to create DCM and established a baseline for the expected growth kinetics in these media. Each *Dysgonomonas* isolate was grown in DMM broth, washed, diluted and subcultured into either DMM or DCM broth and grown to stationary phase. These pre-growth cultures were again washed and diluted into the same or opposite media and incubated aerobically and anaerobically. Aerobically, as expected, all isolates exhibited faster growth rates and higher yield when grown in DCM than in DMM, regardless of the pre-growth medium used (**Figure 6**). Under anaerobic conditions, isolates also achieved greater final yield when grown in DCM rather than DMM, though there was variability between isolates regarding which pre-growth medium allowed the greater yield. These results are unsurprising given reduction in biosynthetic costs of amino acids, vitamins and cofactors provided by the amendments in DCM. Isolates pre-grown in DMM and switched to DCM performed near identically to DCM pre-grown cultures in all cases under aerobic conditions, displaying identical yields and only slight differences in growth rates (**Table 1**). During anaerobic growth there was again variability between isolates with respect to which pre-growth medium resulted in greater final yield. Aerobic cultures pre-grown in DCM and switched to DMM exhibited slightly greater growth rates for all isolates except for GY75, for which DMM-conditioned cells displayed the greater rate. Final yields in DMM during anaerobic growth differed based on pre-growth medium between the isolates, with isolate BGC7 and isolate GY75 displaying greater yield when pre-growth occurred in DCM, while isolate HGC4 and GY617 displayed greater yield when pre-growth occurred in DMM. Perhaps unexpectedly, the switch from DCM to DMM did not elicit an observable lag phase during aerobic growth, which suggested that gene expression related to growth and cell division can shift rapidly following transfer between the two media. Within media types, carrying capacities of the media were highly consistent across isolates and allowed stable population kinetics during stationary phase to at least 96 hours.

### Growth on agar-solidified media

Growth of *Dysgonomonas* isolates on D<u>D</u>M or D<u>M</u>M solidified with 1.5% w/v agar resulted in delayed and overall weak growth when incubated aerobically, while plates incubated anaerobically exhibited good colony growth (author observation). Due to differences in exposure to ambient atmosphere between cultures grown on solid media versus liquid culture, oxidative stress was suspected as a growth-limiting factor on our solid, defined medium. As described by Dione et al. [79], we amended agar-solidified DDM with a single antioxidant (ascorbic acid, uric acid or glutathione) and adjusted pH to 7.5. Isolates were streaked onto each medium, loosely bagged and incubated aerobically. Each antioxidant alone was sufficient to permit aerobic growth on agar-solidified DDM (**Figure 7**). Due to the characteristics of aqueous solutions of ascorbic and uric acid, media required additional pH amendments using concentrated NaOH or HCl. Both ascorbic acid and uric acid changed the hue of the medium to an orange or green-grey color, respectively, which may be inhibitory for some biochemical or genetic analyses. Additionally, the presence of uric acid caused colonies from some isolates to exhibit umbonate morphology (**Figure 7, insets**). Glutathione did not contribute to visible changes in the media or to alterations in colony morphology, and was selected for final media formulations. Addition of ascorbic acid, glutathione or uric acid to liquid media did not significantly alter growth phenotypes in liquid media, with the exception that ascorbic acid lowered growth rate and yield for isolate BGC7 (not shown). Additionally, the *Dysgonomonas* isolates tested did not utilize L-cysteine, ascorbic acid, uric acid or glutathione as a sole source of carbon in DMM (not shown).

To create a parallel complex solid medium, agar-solidified DCM was also amended with glutathione and incubated loosely bagged under common laboratory conditions. Expected growth phenotypes were observed from all isolates on solid DCM after 3 days of incubation under all conditions with the exception that isolates BGC7 and GY75 grew somewhat slower aerobically at 22°C than did the other isolates and required an additional day of incubation (**Figure S8**). Importantly, the creation of DCM, particularly when solidified in the presence of an antioxidant, abolishes the requirement for animal blood when growing *Dysgonomonas*, which may be a limiting factor for some research groups.

When incubated under aerobic conditions, cultures on solid DMM containing glutathione still exhibited unimpressive growth compared to that on DDM, and we speculated that the presence of excess B-vitamins could have contributed to maintaining a reduced intracellular environment during growth on solid DDM. To determine if additional protection from oxidative stress was required, we grew aerobic cultures on solid DMM containing glutathione and incubated plates aerobically, with either unrestricted access to ambient atmosphere (un-bagged plates) or with restricted access to ambient atmosphere (sealed tightly in a plastic bag) (**Figure S9**). Colonies from plates that were un-bagged displayed sparse growth with small colonies which never reached the size of those grown on DDM under similar conditions. In contrast, colonies from plates that were incubated in sealed plastic bags exhibited the typical colony phenotype observed on DDM. We suspect that restricting access to ambient atmosphere could provide several advantages for growth of *Dysgonomonas*; (i) moisture from the media is retained during extended incubation; (ii) oxygen is likely consumed during aerobic respiration faster than it can be replaced in a sealed plastic bag, creating microaerophilic conditions with reduced oxidative pressure; (iii) the buildup of volatile metabolic end products such as carbon dioxide may provide beneficial conditions for growth, as has been demonstrated for several *Dysgonomonas* spp. [9].

### Enrichment of *Dysgonomonas* isolates from termite hindgut

In addition to providing access to ‘cleaner’ physiological and genetics studies, we were interested to determine if DMM could be used to selectively enrich for *Dysgonomonas* from environmental samples. We collected worker and alate termites from the same colony during a swarming event and performed bacterial strain isolation, purification, high-throughput V4 16S rRNA gene screening and ASV generation as described in Materials and Methods. Thirteen isolates representative of four ASV-groups (**Table S4**) were selected for full-length 16S rRNA genes sequencing alongside isolates BGC7, HGC4, GY75 and GY617. We reconstructed a 16S rRNA gene maximum-likelihood phylogeny containing genes from our own isolates and those from other cultured members of *Dysgonomonas* and overlaid ASV-group and isolation source metadata (**Figure 8**). Our small collection of *Dysgonomonas* isolates displayed congruency between ASV groups and full length 16S rRNA gene sequences; that is, all full-length 16S gene sequences from our isolates were also clustered by ASV group. Representative genes obtained from sequence databases that shared 100% identity to, and over the entire length of, our ASVs were considered part of an ASV-group, even if there was disparity between full-length sequences. Sequences in ASV-group 3* represent a subset of 16S rRNA genes from species which were not isolated or used in this study but are identical over the ∼250 bp of the V4 region of 16S rRNA gene that is used to generate ASVs. ASV-group 5 contained only the sequence derived from isolate BGC7, we did not obtain any novel isolates with this ASV during this work.

*Dysgonomonas* isolates that belonged to ASV-group 1 also shared 100% identity between full-length 16S rRNA gene, which included the type strain *D. gadei* (NR_113134.1). Interestingly, we obtained ASV-group 1 isolates from worker and alate hindguts that shared 100% identity with GY75, which was isolated 21 months previous from the same colony. ASVs from isolates within ASV-group 4 shared 98% identity to that of ASV-group 1 (5 substitutions) and clustered separately with *Dysgonomonas termitidis* (AB971823.1) to the exclusion of ASV-group 1 members, based on full-length 16S gene sequences. Full-length sequences belonging to ASV-group 4 shared 98.3% identity with *D. termitidis* and 98.0% identity to those sequences comprising ASV-group 1, representing what could be a novel species (**Files S1 & S2**). Isolates that belonged to ASV-group 2 contained isolate HGC4, isolate GY617, five additional isolates from worker and alate hindguts (WAn2, WAn3, AAn4, AAn7 & AAn9), sp. zg-930 (MN933917.1) and *D. alginatilytica* (NR_137388.1), although *D. alginatilytica* clustered separately based on full-length 16S gene sequence. All termite-derived sequences within ASV-group 2 from this work share >99.9% identity, exhibiting only 1-2 bp substitutions along the full-length 16S gene and together shared >99.9% identity to isolate HGC4, which was isolated from *R. flavipes* termites by a separate laboratory several years prior to our study. Moreover, isolates WAn2 and WAn3 showed 100% identity along the full-length 16S rRNA gene sequence with isolate GY617, which was isolated twenty-one months prior at the same timepoint and from the same termite colony as GY75.

That isolates belonging to ASV-groups 1 and 2 were obtained from worker hindguts from the same colony over multiple seasons, and that isolates belonging to both of these ASV-groups are also able to be obtained from actively-swarming alate hindguts (also from the same termite colony), suggested that these particular strains of *Dysgonomonas* may be stable members of the termite hindgut community within this colony. This is supported by ASV analysis of data from [47] (PRJEB5527) and [100] (PRJEB20463) (conducted using the same methods as described in Materials & Methods) which contained ASVs from termite hindguts within geologically distinct locations across New England which were identical to those found in ASV-groups 1, 2 and 4 from our study (data not shown). Additionally, recent work from our lab [101] which used hindgut DNA from worker termites trapped from the same colony as used in this study, identified Bacteroidetes_ASV008, which was associated with several species of eukaryotic protists, to be identical to the independently generated ASV_4 from this study. Although the diversity of *Dysgonomonas* ASVs in sequencing data from termites suggests many strains are likely acquired horizontally from the environment, our finding that isolates from ASV-groups 1, 2 and 4 contain members isolated from both workers and alates may point to the possibility that some strains of *Dysgonomonas* could gain a competitive advantage in the termite hindgut by being vertically transmitted to new founder populations via alates.

Although studies have shown that high-throughput sequencing of full-length 16S rRNA genes can provide within-species resolution [102], here we have only provided representative sequences pertaining to dominant clones from 16S rRNA gene libraries for each organism. As such, we also do not intend our phylogenetic reconstruction to delimit species or strains. Additionally, we found that isolates which shared ASVs (and in the case of *D. gadei*, 100% similarity over >1.4kb fragments of the 16S rRNA gene) were found in vastly different environments. For example, ASV-group 1 contains sequences from isolates obtained from termite and human sources; ASV-group 3* contains isolates from sludge, microbial fuel cells, aquatic and human sources. It would simple, but incorrect, to imply that ASV groups were correlated with natural reservoir, genomic content or metabolic function, and we do not intend to convey this point.

The use of solid DMM to culture *Dysgonomonas* isolates distributed widely across a 16S rRNA gene phylogeny demonstrates that our chemically defined media are broadly applicable and are likely suitable and otherwise easily amenable for the growth of most strains of *Dysgonomonas*.

## Conclusion

Although *Dysgonomonas* are considered fastidious organisms, their minimal requirements for growth are quite simple. The termite-hindgut derived and phylogenetically diverse *Dysgonomonas* isolates tested in this work exhibit requirements for L-cysteine, ferric hemin, biotin, thiamine and nearly all required cyanocobalamin. Our isolates exhibit preference for media containing ammonium and sulfate and required additional antioxidants when growing aerobically on solid media.

Robust growth on minimal medium is the cornerstone for many physiological and genetic studies using bacteria, and during preparation of this manuscript we were able to further utilize DMM to screen for naturally occurring nucleotide, amino acid and vitamin auxotrophies, examine resistance phenotypes to antibiotics of interest and observe growth phenotypes using specific animal- and plant-derived carbon sources, including polysaccharides associated with lignocellulose (manuscripts in preparation). The media formulations described here provide robust and reliable growth in complex, defined or minimal variations that limit animal-derived components and laboratory equipment such as anerobic chambers. These media can be used in liquid culture, or agar-solidified under both aerobic or anaerobic conditions. The results from this work provide three different, but parallel, media for the growth of members of genus *Dysgonomonas* and will aid in facilitating further physiological and genetic characterizations to determine their functional roles within particular ecological systems.

## Acknowledgements

We gratefully acknowledge Dr. Joerg Graf and Dr. Michael C. Nelson for providing *Dysgonomonas* spp. BGC7 and HGC4, and Dr. Michael Stephens for assistance with DADA2.

---

**Figure S1.**
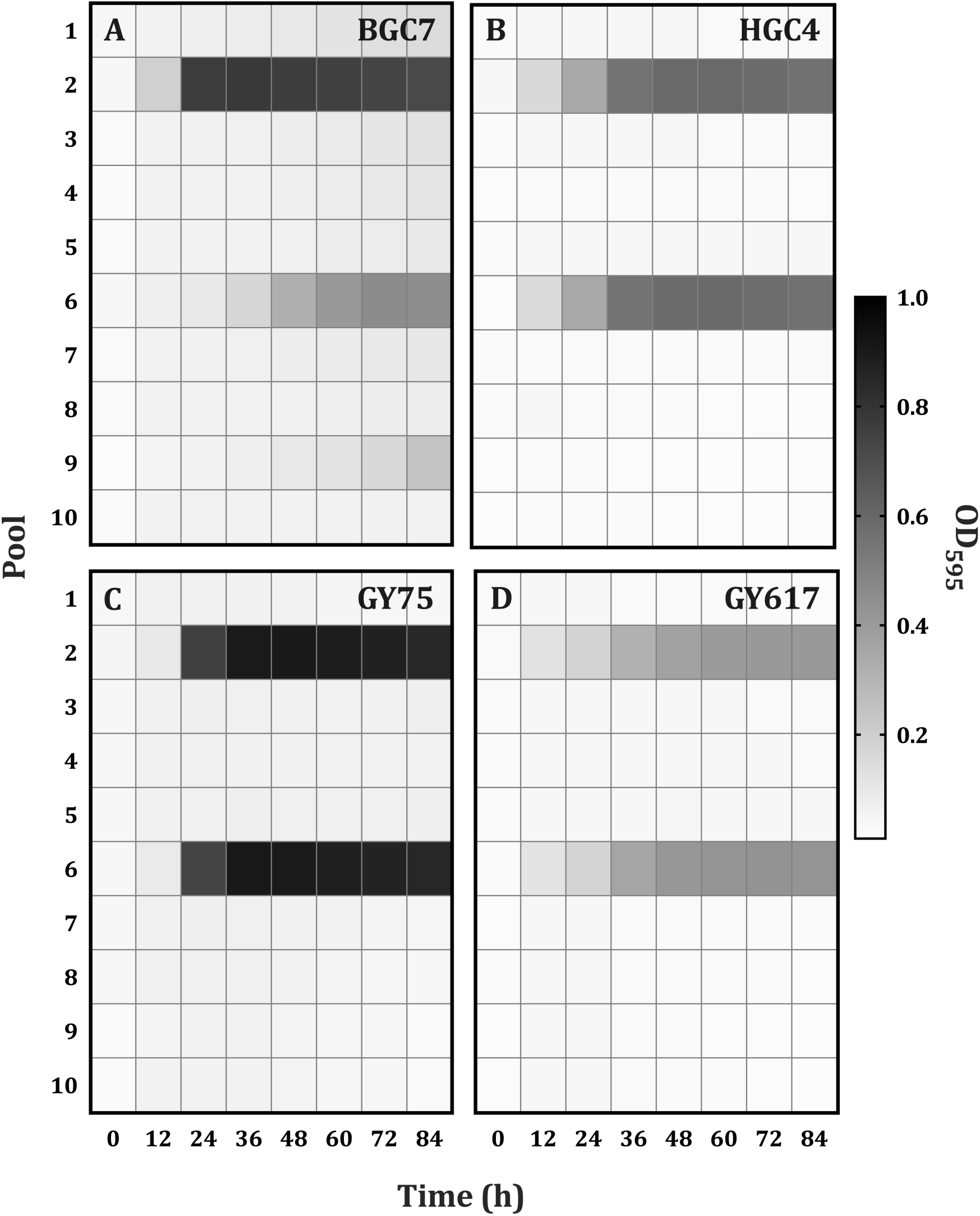
Amino acid auxanography using defined medium. Heat map of corrected growth measurements of *Dysgonomonas* isolates BGC7 (A), HGC4 (B), GY75 (C) or GY617 (D) at twelve-hour intervals. Isolates were grown in 1X M9 salts supplemented with 5% v/v hemin solution, 5% v/v WVS, 0.5% v/v WMS, 0.5% w/v D-glucose, 50 μg/ml kanamycin sulfate, pH 7.5, amended with a single combinatorial amino acid pool, with each amino acid at 0.33 mg/ml. Amino acid pools are as follows: Pool 1: L-phenylalanine, L-serine, L-tryptophan, L-tyrosine, L-glutamine; Pool 2: L-alanine, L-cysteine, L-threonine, L-asparagine, L-methionine, DAP; Pool 3: L-arginine, L-ornithine, L-aspartic acid, L-proline, L-glutamic acid; Pool 4: L-leucine, glycine, L-isoleucine, L-histidine, L-lysine, L-valine; Pool 5: L-phenylalanine, L-alanine, L-arginine, L-leucine; Pool 6: L-serine, L-cysteine, L-ornithine, glycine; Pool 7: L-tryptophan, L-threonine, L-aspartic acid, L-isoleucine; Pool 8: L-tyrosine, L-asparagine, L-proline, L-histidine; Pool 9: L-methionine, L-glutamic acid, L-lysine; Pool 10: L-glutamine, DAP, L-valine, and can also be found in **Table S1**.

**Figure S2.**
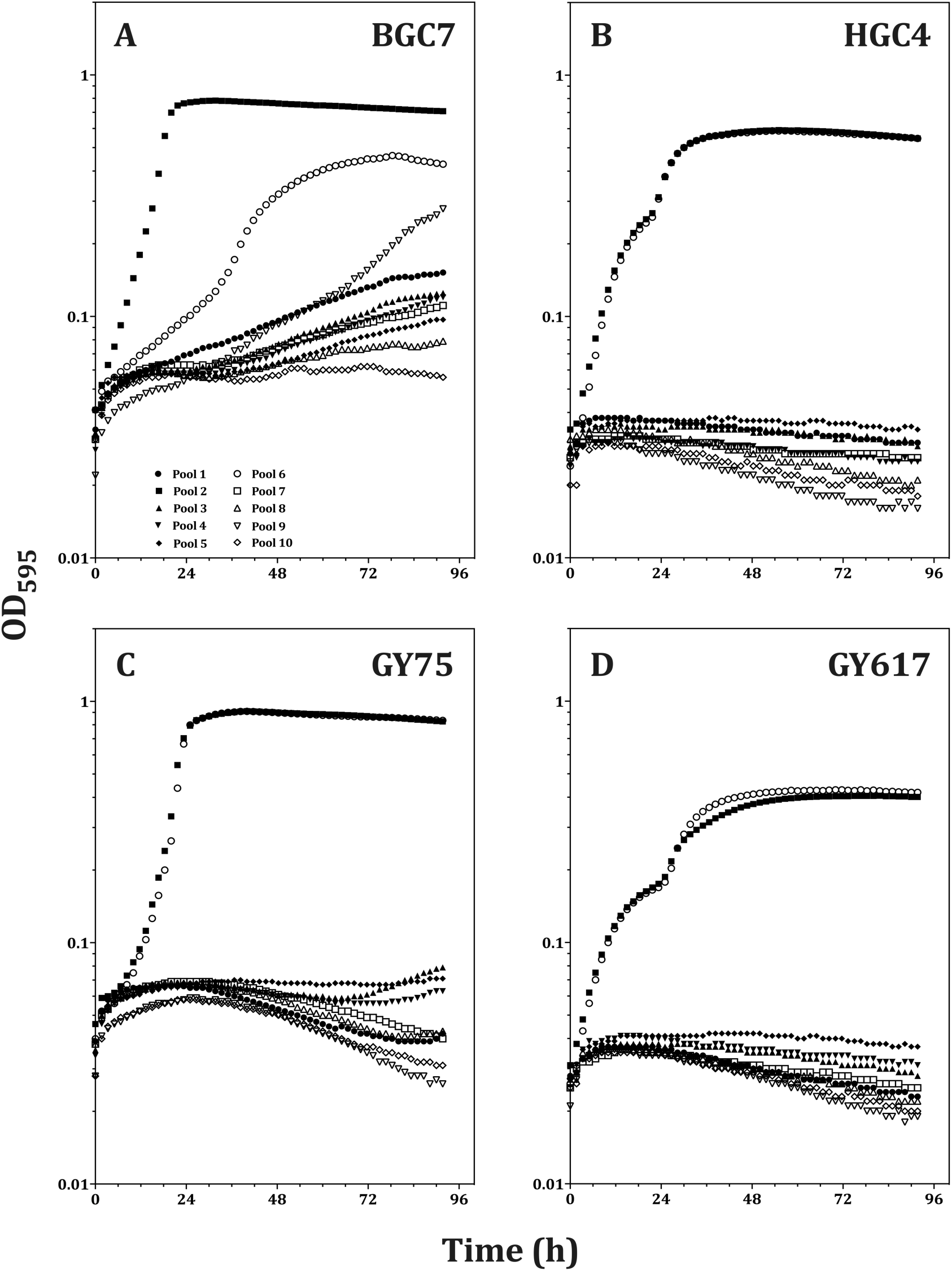
Amino acid auxanography. Growth curves of *Dysgonomonas* isolates BGC7 (A), HGC4 (B), GY75 (C) or GY617 (D) in 1X M9 salts supplemented with 5% v/v hemin solution, 5% v/v WVS, 0.5% v/v WMS, 0.5% w/v D-glucose, 50 μg/ml kanamycin sulfate, pH 7.5, amended with a single combinatorial amino acid pool, with each amino acid at 0.33 mg/ml. Amino acid pools can be found in **Table S1**.

**Figure S3.**
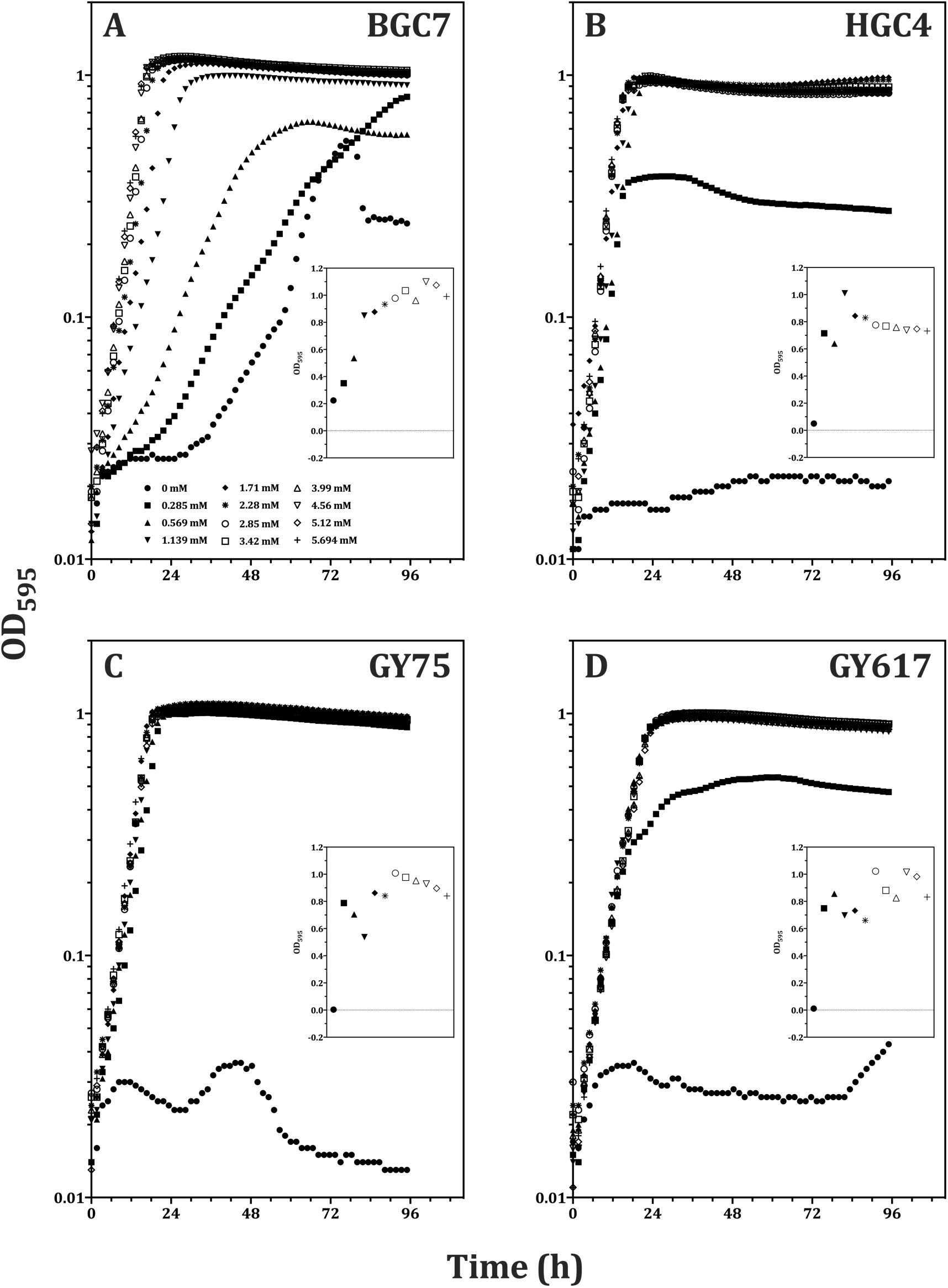
Aerobic growth in defined medium is L-cysteine-dependent. Growth of *Dysgonomonas* isolates BGC7 (A), HGC4 (B), GY75 (C) or GY617 (D) in 1X M9 salts supplemented with 5% v/v hemin solution, 5% v/v WVS, 1% v/v WMS, 0.5% w/v D-glucose, 50 μg/ml kanamycin sulfate, pH 7.5. L-cysteine concentrations are shown in the legend in panel A. Insets show final corrected OD_595_ under anaerobic conditions; the y-axis is in linear units.

**Figure S4.**
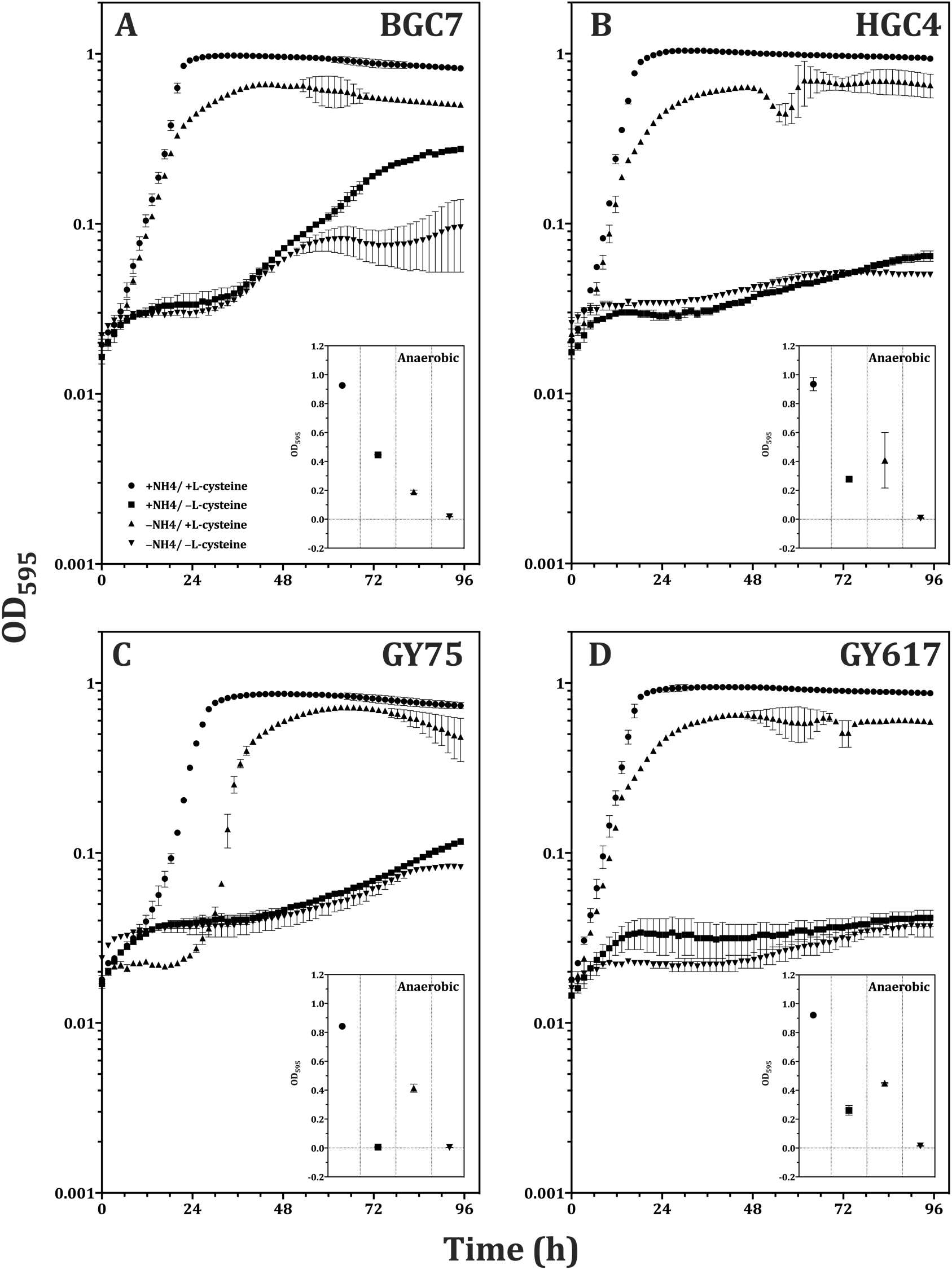
*Dysgonomonas* can utilize L-cysteine as the sole source of nitrogen. *Dysgonomonas* isolates BGC7 (A), HGC4 (B), GY75 (C) or GY617 (D) washed and resuspended in 1X PBS and diluted into 1X M9 salts with or without ammonium chloride, supplemented with 1% v/v WMS, 0.5% w/v D-glucose, 50 μg/ml kanamycin sulfate, pH 7.5, with and without 1.7 mM L-cysteine or L-methionine, as outlined in the legend in panel A. Insets show final corrected OD_595_ under anaerobic conditions; the y-axis is in linear units. Error bars represent standard error of the mean (n=2).

**Figure S5.**
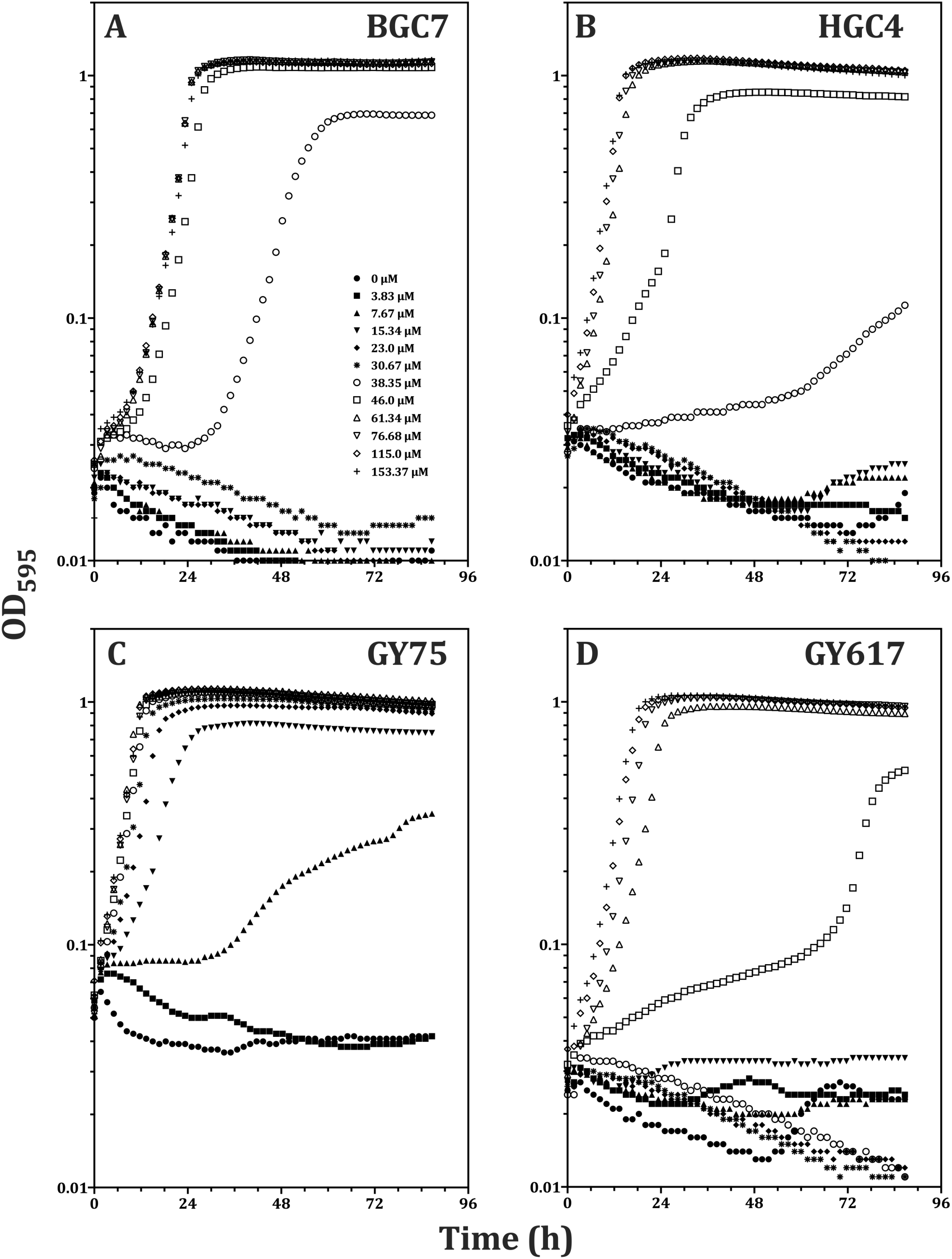
Aerobic growth is dependent on ferric hemin concentration. *Dysgonomonas* isolates BGC7 (A), HGC4 (B), GY75 (C) or GY617 (D) were pre-grown in DDM without hemin, washed and diluted into 1X M9 salts supplemented with 5% v/v WVS, 1% v/v WMS, 0.5% w/v D-glucose, 1.7 mM L-cysteine, 50 μg/ml kanamycin sulfate, pH 7.5. ferric hemin amendments can be found in the legend in panel A.

**Figure S6.**
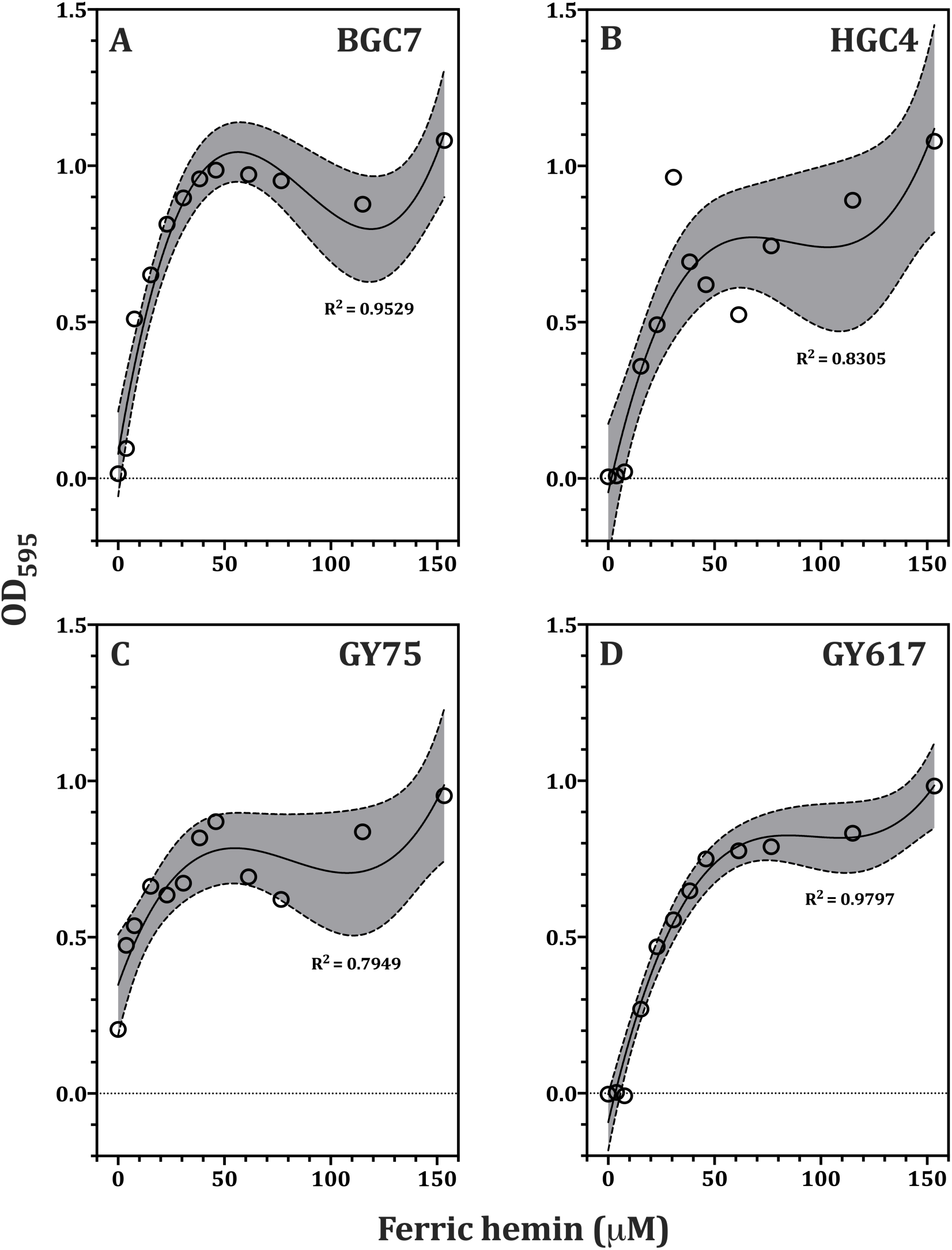
Anaerobic growth yield is dependent on ferric hemin concentration. *Dysgonomonas* isolates BGC7 (A), HGC4 (B), GY75 (C) or GY617 (D) were pre-grown on DDM lacking hemin, washed and diluted into 1X M9 salts supplemented with 5% v/v WVS, 1% v/v WMS, 0.5% w/v D-glucose, 1.7 mM L-cysteine, 50 μg/ml kanamycin sulfate, pH 7.5, and incubated anaerobically. Shown are the final growth yields (difference in OD_595_ at *t*=0h and *t*=85h) across a range of hemin concentrations during anaerobic growth. Identical concentrations as in Fig. 5 were used to determine final yield. The data were fit using third order polynomial regression (solid line) and coefficients of determination for the fit are given as R^2^ values as indicated. Confidence intervals of 95% are represented by shading bounded by dashed lines.

**Figure S7.**
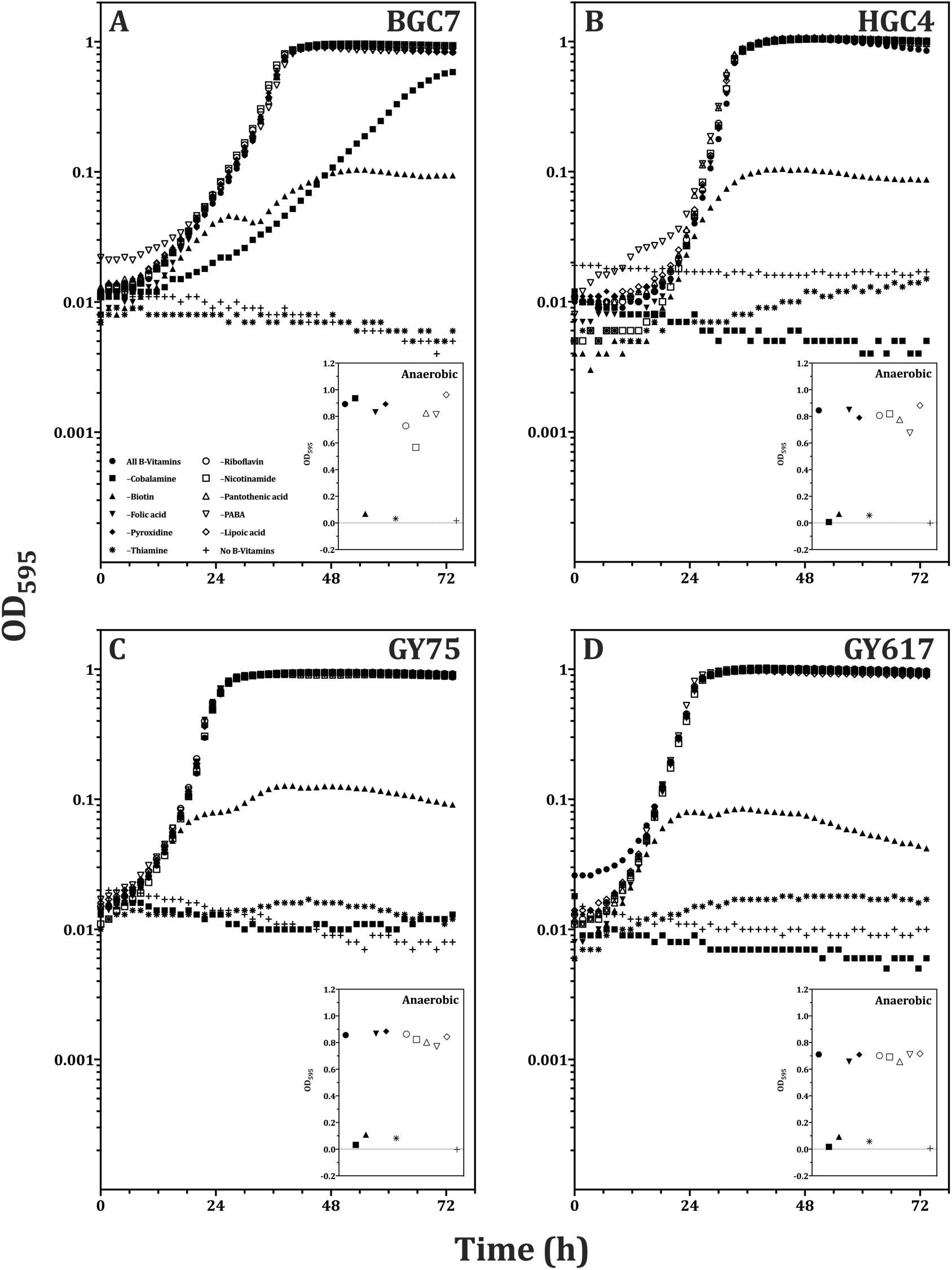
Isolates exhibit multiple B-vitamin auxotrophies. *Dysgonomonas* isolates BGC7 (A), HGC4 (B), GY75 (C) or GY617 (D) were pre-grown in DDM lacking B-vitamins, washed and diluted into 1X M9 salts supplemented with 1% v/v WMS, 0.5% w/v D-glucose, 1.7 mM L-cysteine, 10% v/v hemin solution, 50 μg/ml kanamycin sulfate, pH 7.5, containing vitamin pools with all, none or a single cofactor omission, as outlined in the legend in panel A. Insets show final corrected OD_595_ under anaerobic conditions; the y-axis is in linear units.

**Figure S8.**
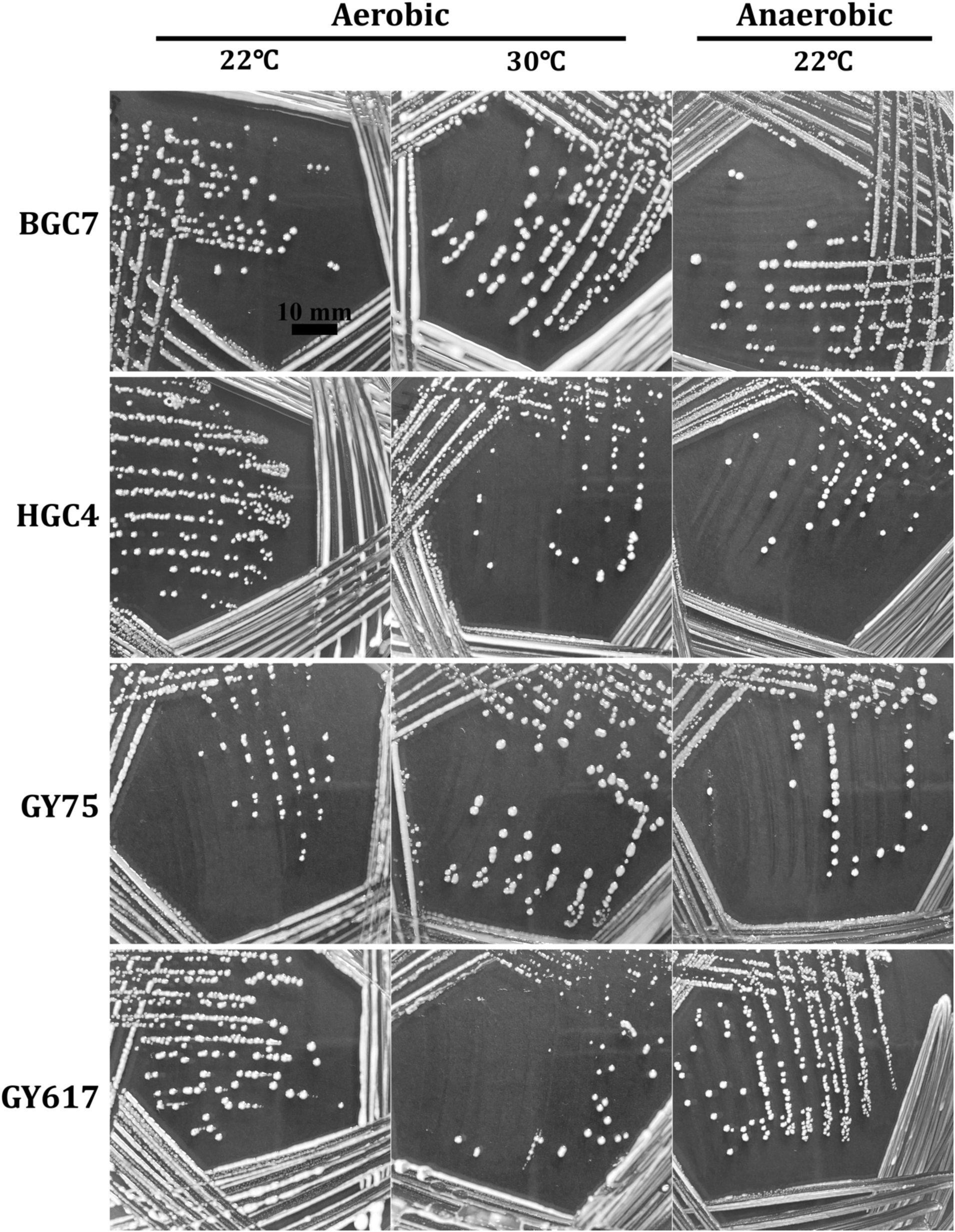
Agar-solidified complex medium (DCM) supports growth under common incubation conditions. *Dysgonomonas* isolates were streaked from solid DDM onto agar-solidified DCM containing 0.1 mg/ml reduced glutathione and streaked to isolation. Plates were loosely bagged, incubated aerobically at 22°C or 30°C or anaerobically at 22°C for 4 days and photographed. Scale bar represents 10 mm.

**Figure S9.**
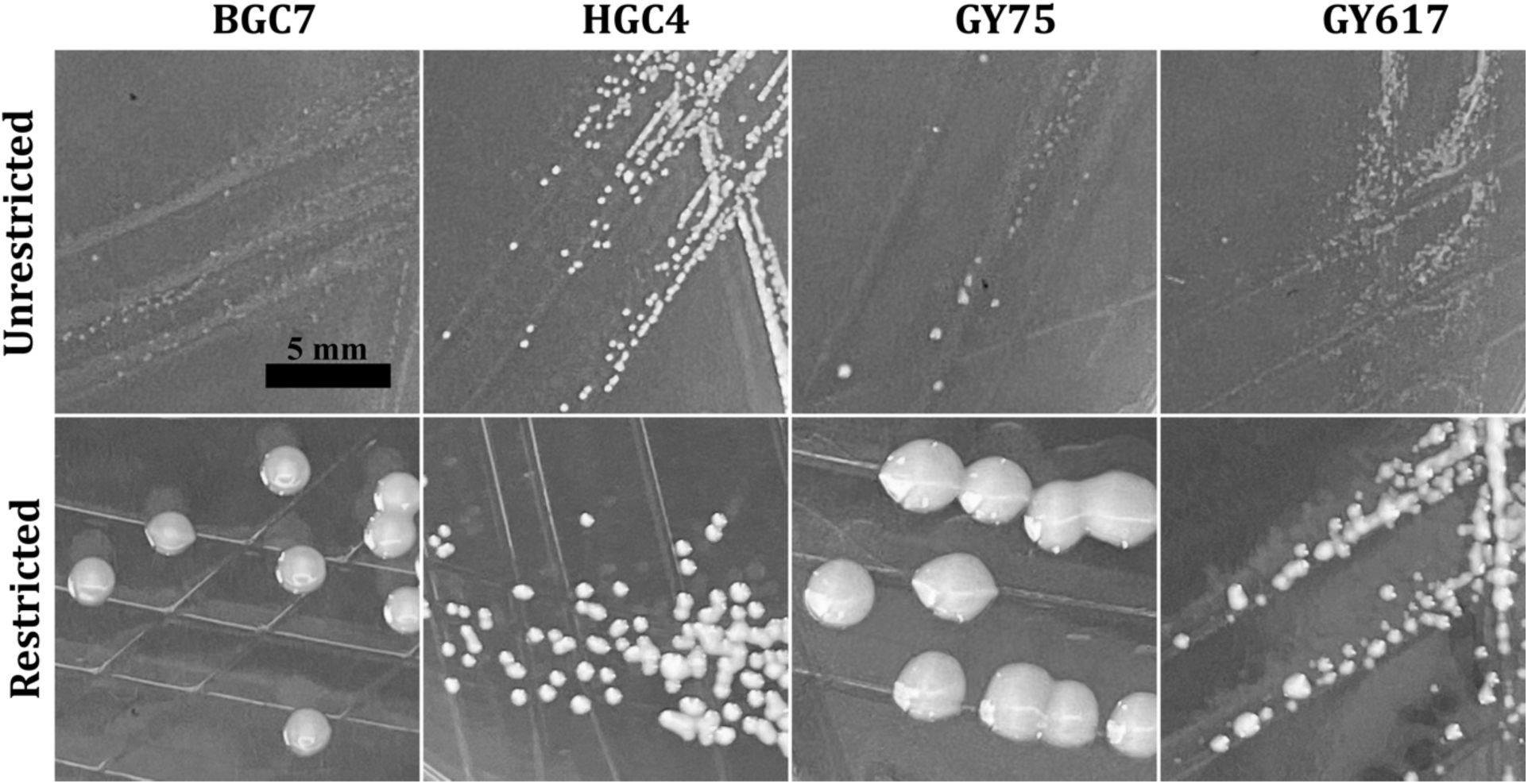
Incubation in atmosphere-restricted conditions promotes growth on agar-solidified DMM containing antioxidants. Several colonies of each *Dysgonomonas* isolate were taken from solid DDM and separately pooled in M9 salts, spotted and streaked onto agar-solidified DMM containing 0.1 mg/ml glutathione. Plates were incubated at 30°C with either unrestricted access (un-bagged) or restricted access (sealed in plastic bag) to ambient atmosphere for 5 days and photographed. Scale bar represents 5 mm.

**Table S1. Amino acid auxanography pools.** Twenty-two amino acids were prepared as aqueous 10 mg/ml solutions. Ten pools were constructed such that concentrations of each component was 1.67 mg/ml. Pools were added to media such that final concentration of each amino acid was 0.33 mg/ml.

**Table S2. Recipes for stock salt, metal and vitamin solutions.** Recipe for concentrated stock solutions are listed in mass per liter and molarity. Stock solutions can be prepared and safely stored at 4°C for weeks to months with minimal effect on growth. See references for WMS and WVS in Materials and Methods for preparation instructions.

**Table S3. Media recipes for DCM, DDM & DMM.** Media recipes for DMM, DDM and DCM are provided along with final concentrations in mass per liter and molarities. Suggestions for preparation are also provided, but we assume some knowledge and experience with media preparation.

**Table S4. ASVs generated from *Dysgonomonas* isolates used in this study.** Thirty isolates were obtained from worker and alate termites and purified on DMM. The V4 region of the 16S rRNA gene was amplified and sequenced using dual-barcoded primers on an Illumina MiSeq platform. The R package DADA2 was used to generate ASVs. Molecular-grade water was used as a no-template control, and *Dysgonomonas* spp. BGC7 and HGC4 as well as isolates GY75 and GY617 were used as biological controls. Read counts less than 5 percent of the total attributed to a particular isolate were considered amplification or sequencing errors that passed filtering and disregarded. ASV_3* was not generated from high-throughput sequencing data, but was observed to represent a subset of dideoxy-sequenced 16S rRNA genes which share 100% similarity over the same V4 region as ASVs generated with DADA2 (see **Figure 13**).

**File S1. Multiple sequence alignment of 16S rRNA genes used in this study (FASTA format).**

**File S2. Pairwise nucleotide identities for 16S rRNA genes used in this study (CSV format).**

